# SOP-Multi-*π*: A Two-Site Coarse-Grained Model with Rescaled Aromatic Interactions to Bridge Single-Chain Ensembles and Condensate Phase Behavior of IDPs and Multidomain Proteins

**DOI:** 10.64898/2026.09.30.755554

**Authors:** Santosh Prajapati, Neha Mathur, Debayan Chakraborty, Anand Srivastava

## Abstract

We present SOP-Multi-*π*, an enhanced version of the two-bead-per-amino-acid resolution Self-Organized Polymer (SOP) model designed to investigate the conformational dynamics of Intrinsically Disordered Proteins (IDPs) and their phase behavior in biomolecular condensates. The original SOP-IDP and SOP-Multi models work well for chains in good solvents that predominantly adopt expanded conformations. However, they fail to reproduce the single-chain properties and phase-separation behavior of IDPs such as hnRNPA1 and its variants, which have a distinct pattern of aromatic residues. We therefore reparameterized the SOP-IDP force field to explicitly capture *π*–*π* and cation–*π* interactions at the coarse-grained (CG) resolution. These interactions are weak but important for biomolecular recognition and for driving and stabilizing biomolecular condensates. Dimensionless rescaling prefactors (*λ_ij_*) were applied exclusively to the aromatic *π*–*π* (Phe, Tyr, Trp) and cation–*π* (Arg/Lys with Phe, Tyr, Trp) pairs. They were optimized iteratively against experimental radii of gyration of the hnRNPA1 low-complexity domain (A1-LCD) and its 16 aromatic and charge mutants. This restored a physically meaningful Tyr *>* Phe interaction hierarchy, and the Pearson correlation for *R_g_* improved from *r* = 0.07 (SOP-IDP) to *r* = 0.83 (RMSD = 0.08 nm). The same parameters, without further tuning, gave *r* = 0.93 for 34 additional IDPs (24–198 residues) and reproduced Flory scaling (*ν* ≈ 0.588). We back-mapped microsecond-scale CG ensembles to all-atom representations and computed SAXS and Kratky profiles for direct comparison with experiments on IDPs and on the multidomain proteins hnRNPA1, TDP-43, G3BP1, and FUS. In direct-coexistence slab simulations, the model captures the phase behavior of A1-LCD condensates and their variants, with critical temperatures that correlate strongly with experiment (*r* = 0.88) but are somewhat underestimated in absolute terms. The model is implemented in LAMMPS, and a detailed user manual is provided for setting up single-chain and multi-chain condensate simulations of full-length multidomain proteins.

## 1 Introduction

Biomolecular condensates (BMC) formed via liquid-liquid phase separation (LLPS) of intrinsically disordered proteins (IDPs) and regions (IDRs) have emerged as pivotal organizers of intracellular spatiotemporal compartmentalization, driving processes ranging from ribogenesis and RNA processing to signal transduction and transcriptional control.^1,2^ Within the theoretical framework of polymer physics, the phase behavior of these multivalent biopolymers is primarily dictated by a “sticker-and-spacer” architecture, wherein short-range, non-covalent physical cross-links act as associative “stickers” interspersed among flexible, solvated “spacers”.^3,4^ Among the diverse set of weak non-covalent forces underpinning condensate cohesion, *π*-associated interactions, specifically *π*-*π* aromatic stacking and cation-*π* contacts (most prominently between the Guanidinium group of Arginine and the aromatic rings of Tyrosine or Phenylalanine) serve as indispensable molecular drivers of condensation. ^3,5–7^ Crucially, the thermodynamic driving force for LLPS, characterized by the saturation concentration (*c*_sat_) and upper critical solution temperature (UCST) phase boundary, depends not merely on the total valence (number) of these interactive sticker residues, but explicitly on their specific spatial sequence patterning along the primary polypeptide chain.^8–10^ Functionally, precise evolutionary tuning of valence and sequence patterning allows cells to maintain dynamic, reversible membraneless organelles that rapidly assemble and dissolve in response to microenvironmental cues or post-translational modifications. Conversely, disease-associated missense mutations or aberrant post-translational hypocondensation events that alter cation–*π* balance or enhance local aromatic valence frequently perturb this delicate molecular grammar, shifting the phase equilibrium away from dynamic liquid-like droplets toward irreversible, aberrant liquid-to-solid phase transitions-promoting the nucleation of cytotoxic fibrillar aggregates and pathological amyloid hydrogels implicated in neurodegenerative disorders including amyloid beta (A*β*) and tau aggregation in Alzheimer’s disease (AD), intracellular *α*-synuclein aggregates in Parkinson’s disease (PD), polyQ-driven protein aggregates in Huntington’s disease (HD), and TDP-43 and FET protein aggregates in amyotrophic lateral sclerosis (ALS) and frontotemporal dementia (FTD).^11,12^

Recently, Tanja Mittag and Rohit Pappu groups used the low-complexity domain of hnRNPA1 (A1-LCD) as a strikingly clean model system to isolate how the molecular grammar of aromatic residues based “sticker” interactions (cation – *π* and *π* – *π*) govern phase separation with carefully designed mutant panels. In the 2020 Science paper, ^9^ the authors built a graded series of A1-LCD variants that systematically titrated the number of aromatic (Tyr/Phe) residues while holding overall composition constant—constructs named, for instance, A1-LCD Aro-, Aro–, and Aro+, such that valence (the number of stickers) could be decoupled from patterning (their spacing along the chain). This allowed them to show, using an integrated simulation-experiment-theory approach, that the extent of chain compaction and the driving force for phase separation are directly determined by aromatic valence, and that natural sequences favor distributed rather than clustered aromatic residues, with disruption of this patterning promoting irreversible aggregation instead of reversible liquid-liquid phase separation. The 2022 Nature Chemistry follow-up work^10^ extended this logic to the interplay between aromatic and charged residues by curating a second, equally deliberate and elegant panel of mutant systems —exemplified by variants such as A1-LCD-12F+12Y, in which twelve Phe residues were swapped for Tyr, and A1-LCD+7F-7Y, in which seven Tyr residues were swapped for Phe — which cleverly holds total aromatic valence fixed while changing only the chemical identity (Tyr vs. Phe) and hence the strength of *π* -*π* versus cation *π* coupling with neighboring Arg residues, while other constructs reshuffled the spacing between aromatics and charged residues to probe evolutionarily conserved sequence “grammar”. Together, this pair of meticulously matched, minimally perturbed variant libraries (see Fig. 1) let the researchers cleanly attribute changes in saturation concentration and condensate stability to the valence, chemical identity, and spatial patterning of residues mediationg *π*-*π* and cation *π* interactions, rather than to confounding changes in net charge, hydrophobicity, or chain length—making these among the most rigorously controlled sequence-to-phase-behavior studies in the intrinsically disordered protein field.

**Figure 1:**
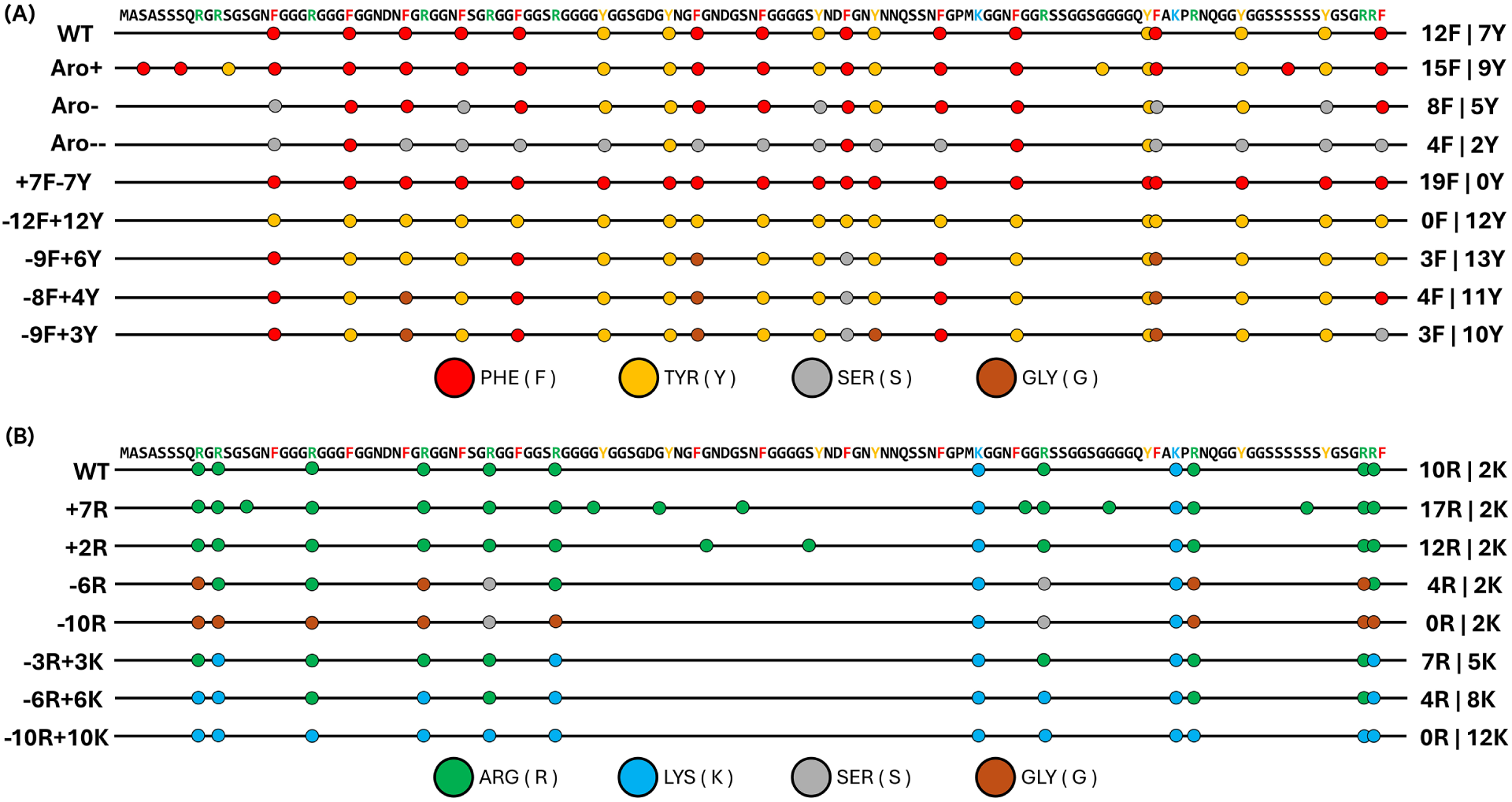
(A) Representing the positioning of aromatic residues (Phe and Try indicated as red and yellow circles, respectively, and Gly and Ser indicated as grey and brown circles, respectively) along the WT(A1-LCD) sequence and its variants.^9,10^ The number of Phe(F) and Tyr (Y) is indicated on the right-hand side. (B) Representing the positioning of positively charged residues (Arg and Lys indicated as green and skyblue circles) along the WT(A1-LCD) sequence and its variants.^10^ The number of Arg(R) and Lys (K) is indicated on the right-hand side.

**Figure 2:**
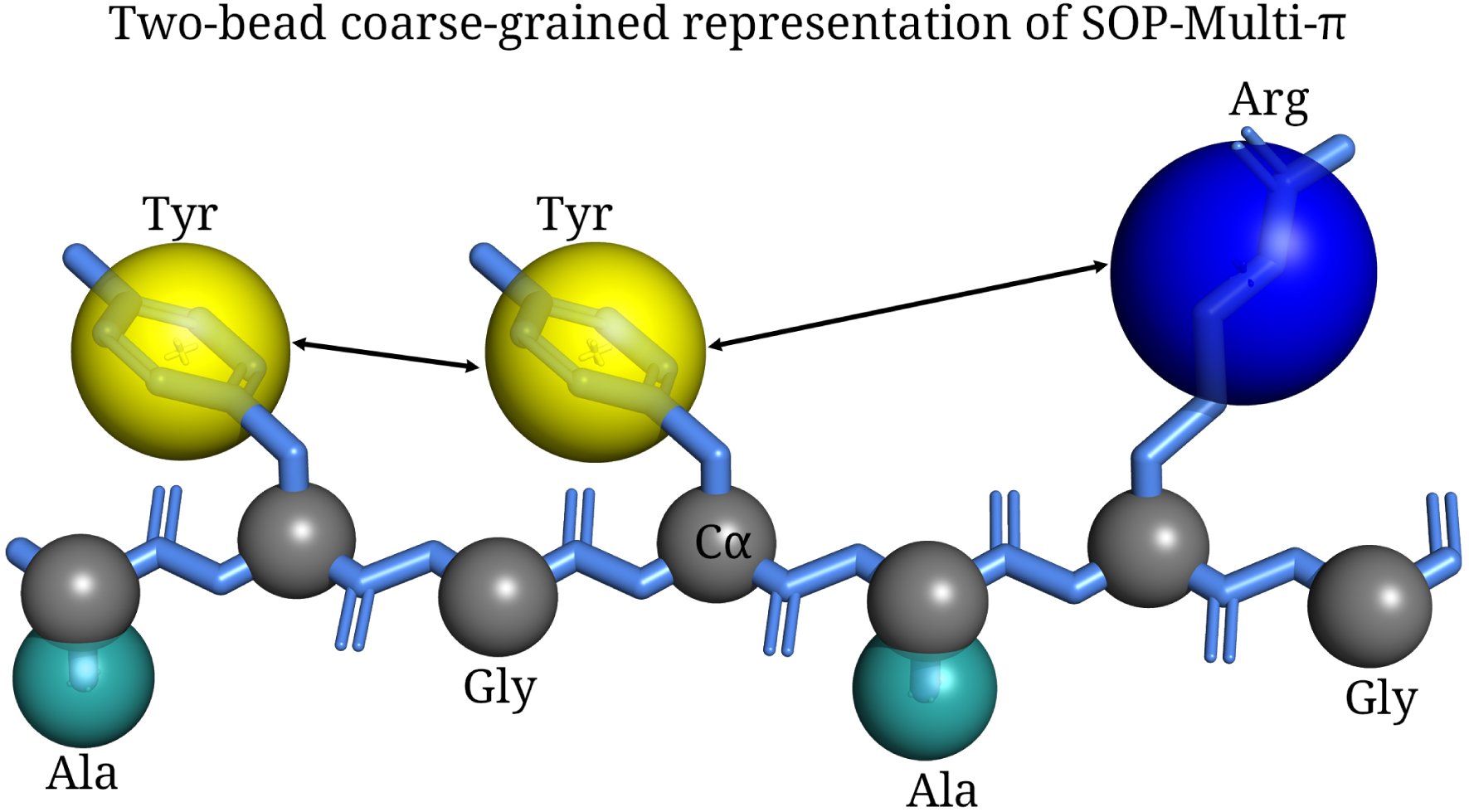
Schematic diagram of a two-bead coarse-grained model representation of a 7-amino acid peptide. The arrows indicate *π* − *π* and cation-*π* interactions.

Biophysical experimental methodologies have illuminated the structural ensembles and conformational dynamics of IDPs both in the dilute solution phase as well as inside condensate phases. However, while these experimental techniques provide indispensable macroscopic and ensemble-averaged readouts, they usually lack the spatiotemporal resolution needed to delineate the precise menagerie of dynamic and transient but important residue-residue interactions within biomolecular condensates (BMCs).^13^ In this context, Physics-grounded molecular simulations have emerged as a potent tool to elucidate the molecular driving forces underlying the conformational ensembles of IDPs and the formation of biomolecular condensates. Over the past decade (2015-2026), advances in biomolecular simulations—spanning all-atom molecular dynamics (AAMD) to residue-level coarse-grained (CG) models have shed transformative light on the inner workings of IDP conformations and LLPS..^12,14–16^ Nevertheless, the biomolecular simulation methods face distinct multiscale trade-offs and limitations. High-resolution atomistic models require prohibitively high computational resources, rendering them incapable of simulating the system sizes and timescales necessary to observe macroscopic condensate assembly or to calculate full coexistence phase boundaries in a high-throughput manner. Conversely, while residue-level coarse-grained models (typically 1-site or 2-site per amino acid) easily bridge these length and time scales, standard CG models have historically struggled to capture complex, highly directional, or electronic interactions. Single-bead representations often fail to accurately model the anisotropic geometry and multibody quantum chemistry of *π*-*π* aromatic stacking and cation-*π* interactions with notable exceptions like the Mpipi force field.^17^ Other reparameterized models^18,19^ often explicitly reweight or incorporate directionally tuned short-range potentials to recapitulate experimental readouts using a top-down fitting approach. Highly coarse-grained architectures often lack the physical sophistication required to recapitulate subtle cation-*π* and *π*-*π* directional interactions without sacrificing computational efficiency, highlighting a pressing need for next-generation multiscale force fields tailored for IDP phase behavior.

To address this critical gap in coarse-grained methodologies, our group has extended the Self-Organized Polymer (SOP) modeling framework to explicitly capture the complex balance of short-range anisotropic forces in intrinsically disordered systems. Building upon the SOP-IDP architecture by Thirumalai and coworkers, which successfully captured Flory scaling laws, sequence-dependent chain dimensions, and structural heterogeneity of isolated IDPs,.^20,21^ Our group recently developed the 2-site-per-amino-acid SOP-MULTI model; ^22^ here, we present an advanced and highly transferable coarse-grained force field (SOP-Multi-*π*). SOP-MULTI established an integrative platform capable of simultaneously sampling folded domains and disordered linkers in full-length multidomain proteins (such as FUS, G3BP1, hnRNPA1, TDP-43, and TIA1) consistent with small-angle X-ray scattering (SAXS) data. In this advanced version of the SOP-Multi model, we systematically improve the short-range potential energy terms to accurately represent cation-*π* (e.g., Arg-Tyr/Phe) and *π*-*π* aromatic stacking contacts within the 2-site SOP representation. By refining the non-bonded interaction matrix with these experimentally validated *π*-associated parameters, this refined SOP framework successfully bridges single-molecule conformational properties and multi-chain liquid-liquid phase-separation behavior. Specifically, unlike several other CGFF models, SOP-Multi-*π* enables accurate predictions of single-chain conformational ensembles while simultaneously driving quantitatively reliable macroscopic condensate simulations, including phase coexistence boundaries and critical temperature features that the classical SOP-IDP model could not capture. This advancement offers a computationally efficient yet physically rigorous computational platform to dissect the precise molecular grammar governing biomolecular condensation and pathological liquid-to-solid transitions in large-scale multi-protein assemblies.

The rest of the article is organized as follows: the details of the SOP-Multi-*π* model and its force field parameters, along with descriptions of the analysis, are provided in the Materials and Methods section. We report the simulation results alongside experimental data in the Results and Discussion section. Further, we have discussed the scope and limitations of the SOP-Multi-*π*.

## 2 Materials & Methods

### 2.1 SOP-Multi-*π* Model

In the SOP-IDP Model,^20,21^ each amino acid is represented by two beads except glycine. One is a backbone (BB) bead at the center of the *C_α_* carbon atom, and the other is a sidechain (SC) bead located at the center of mass of the sidechain. The van der Waals radius, mass, and charges of each amino acid are tabulated in the supporting information Table S1.

The SOP-IDP potential total energy function is given by:

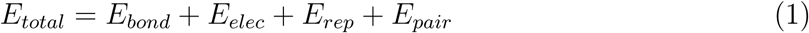

where *E_bond_*represents the bonded interaction, *E_elec_* describes the electrostatic screening interaction among the charged amino acids, *E_rep_*describes the soft repulsive potential, *E_pair_* represents the pair interactions between amino acids.

#### Bonded interaction

The bond energy is described by the finitely extensible non-linear elastic (FENE) potential.

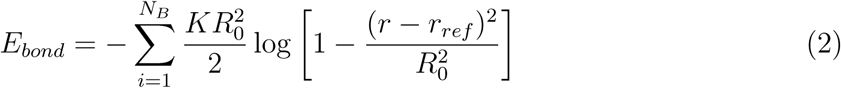

where the spring constant *K* was set to 20 Kcal/mol-Å^2^, *R*_0_ represents the tolerence and was set to 2.0 Å, and the reference bond length for BB-BB beads (*r_ref_*) is 3.8 Å, and for BB-SC beads (*r_ref_*) is the sum of van der Waals radii of BB and SC beads.

#### Electrostatic screening interaction

The salt-screened electrostatic interaction between charged amino acid beads is described using the Debye-Hückel potential.^23^

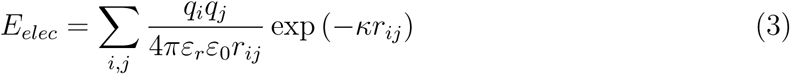

where the *q_i_* and *q_j_* represent the charge on the *i^th^* and *j^th^* beads, and the charge corresponding to the residue is assigned to the sidechain bead. The permittivity of vacuum is *ɛ*_0_. *r_ij_* is the distance between the *i^th^* and *j^th^* beads, 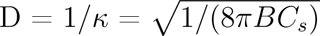 is the Debye screening length of an electrolyte solution of ionic strength *C_s_*, and *B*(*ε_r_*) is the Bjerrum length.^24^ We use the following empirical law^25^ to model the temperature-dependent relative dielectric constant (*ε_r_*) of the implicit aqueous solution. We used a cutoff of 50 Å for electrostatic interaction for single-chain IDPs studies.

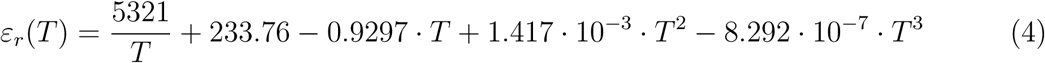

#### Repulsive excluded volume interaction

The repulsive interaction prevents unphysical overlap between the beads and acts between beads separated by a distance of 1-3.

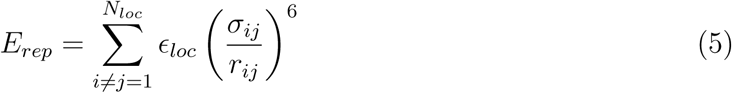

where *ɛ_loc_* is set to 1.0 kcal/mol, *σ_ij_* is sum of the van der Waals radii of beads *i^th^* and *j^th^* and *r_ij_* is distance between the *i^th^*and *j^th^* beads.

#### Pair interaction

Lennard-Jones (LJ) potentials define the non-bonded pair interactions, which are segregated into three types: backbone-backbone (*E_BB_*), backbone-sidechain (*E_BS_*), and sidechain-sidechain (*E_SS_*) interactions,^21^ where the E*_SS_* term encodes the sequence specificity. To reparametrize the *π*−*π* and cation-*π* interactions in the SOP-IDP model, we further divided the *E_SS_* term into rescaled and non-rescaled pairs.

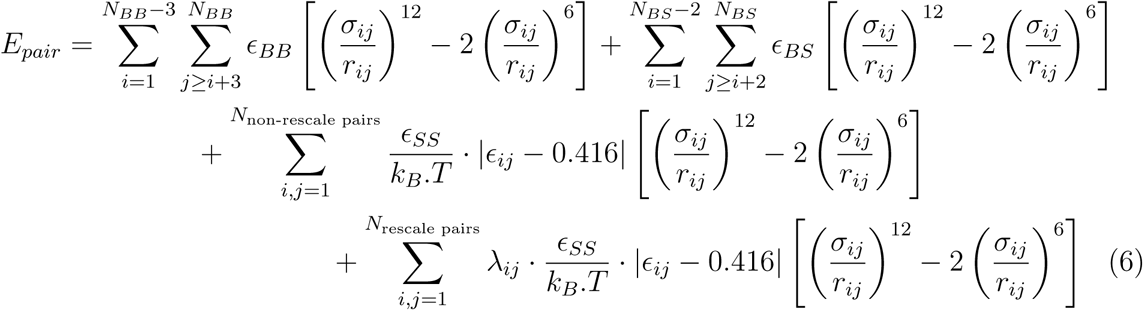

where *ɛ_BB_*, *ɛ_BS_*, and *ɛ_SS_* denote the strength of the interaction between backbone-backbone (BB), backbone-sidechain (BS), and sidechain-sidechain (SS) beads, respectively. Following Mugnai et al.^21^ we set *ɛ_BB_* = 0.104 kcal/mol; *ɛ_BS_* = 0.137 kcal/mol; *ɛ_SS_* = 0.180 kcal/mol. The parameter *ɛ_ij_* is obtained from the Betancourt-Thirumalai statistical potential. ^26^ The *ɛ_ij_* values are unique to each pair of amino acid residues; therefore, they depend explicitly on the IDP sequence. *σ_ij_* is the sum of van der Waals radii of *i^th^* and *j^th^* residues and *r_ij_* is distance between *i^th^* and *j^th^* residues. *k_B_* is the Boltzmann constant, T is the temperature (*k_B_.T* = 0.593 Kcal/mol), and *λ_ij_* is a dimensionless rescaling prefactor that varies across rescaling pairs. In this work, we deduce the *λ_ij_* pairs that capture the subtle *π*-*π* and cation *π* interactions. We used a cutoff of at least 3*σ* for the LJ interaction.^21^

### 2.2 Simulation of the IDPs and multidomain proteins

Each IDP protein was considered as a linear chain embedded in a large cubic box of dimension 50 nm. Each multidomain protein initial configuration was generated according to Baratam et al.^22^ in a cubic box of dimension 50 nm using the AlphaFold^27^ generated PDB file. Molecular dynamics simulations were performed in the canonical ensemble (NVT), using the Langevin thermostat^28^ with a relaxation time of 10 ps and a time step of 10 fs. The simulations were carried out for about 10 *µ*s, keeping the conditions as close as possible to the experimental setup. All the experimental conditions taken according to the article Martin et al.^9^ and Bremer et al. ^10^ All simulations were carried out using the LAMMPS software.^29^

To recalibrate the *π* − *π* and cation−*π* interactions, we selected a total of 16 hnRNPA1-LCD variants as shown in the Fig. 1. We selected IDPs dataset from. Martin *et al.*^9^ (WT and its three variants (Aro+, Aro−, and Aro−−)) and from Bremer *et al.*^10^ (+7F−7Y,−12F+12Y, −9F+6Y, −8F+4Y, and −9F+3Y). All the variants have the same number of charged residues. The schematic illustrating the positioning of aromatic residues along the sequence is shown in Figure 1A. To reparametrize the cation-*π* interaction, we have selected the IDPs dataset from Mittag and co-workers^10^ (+7R, +2R, −6R, −10R, −3R+3K, −6R+6K, and −10R+10K). All the variants have the same number of aromatic residues. The schematic illustrating the positioning of charged residues along the sequence is shown in Fig. 1B. The sequence properties of A1-LCD and its variants are tabulated in the supporting information Table S2.

The differential sticker strengths of Arginine (Arg) and Lysine (Lys) represent one of the most consequential and experimentally well-documented distinctions in the molecular grammar of biomolecular condensate formation. Within the stickers-and-spacers framework, Arg functions as a context-dependent auxiliary sticker. In contrast, Lys not only fails to act as a sticker but actively weakens sticker–sticker interactions. ^3^ This asymmetry is particularly striking, given that both residues carry identical net positive charges at physiological pH. To systematically reparameterize cation−*π* interactions in the SOP-IDP model and capture this Arg-Lys asymmetry, we selected a panel of seven A1-LCD sequence variants from Bremer et al.,^10^ designed to independently modulate the number and identity of positively charged residues while holding the aromatic residue content fixed. The first subset of A1-LCD variants systematically modulates total Arg content. The +7R variant introduces 7 additional Arg residues into the A1-LCD sequence, increasing the total Arg count and thereby amplifying the available cation−*π* contacts with the fixed aromatic sticker array. Conversely, the +2R variant adds only 2 Arg residues, providing an intermediate perturbation. On the depletion side, the −6R variant removes 6 Arg residues from the A1-LCD sequence, substantially reducing cation−*π* interaction capacity, while the −10R variant eliminates 10 Arg residues. The second subset of variants −3R+3K, −6R+6K, and −10R+10K are designed to isolate the qualitative difference in sticker identity between Arg and Lys while maintaining overall charge neutrality. In these mutant systems, 3, 6, and 10 Arg residues are replaced by an equivalent number of Lys residues, respectively, such that the total number of positively charged residues and the net charge of the sequence remain unchanged.^10^ The schematic is shown in Figure 1.

### 2.3 Bimolecular condensate simulation

The phase diagrams of the different systems have been evaluated by means of Direct Coexistence simulations.^30^ The Direct Coexistence simulations allow simultaneous simulation of both the dense and dilute phases of the mixture. Additionally, these simulations are performed with periodic boundary conditions in each dimension. To initiate the direct coexistence simulations, we start with a single copy of IDPs in a cubic box and replicate the box in the x, y, and z directions to obtain the target number of chains (125) in the LAMMPS. ^29^ We then iteratively scaled the simulation box to compress it until a high chain density of ≈ 0.9 − 1 g/cc^3^ was reached. We elongate the box along the z direction, keeping the x and y dimensions fixed.^31^ Here, we use the Langevin thermostat at the target temperature, with a relaxation time of 5 ps and a time step of 10 fs. We have simulated the entire system for about 1 *µ*s. The details of the systems are reported in the Table 1.

**Table 1:** All four systems have 125 chains in the slab box with the box dimension as mentioned below.

| System | chains | Beads | Lx=Ly (nm) | Lz (nm) |
| --- | --- | --- | --- | --- |
| WT | 125 | 27375 | 13.9 | 97.0 |
| +7F-7Y | 125 | 27250 | 13.7 | 96.0 |
| -12F+12Y | 125 | 27250 | 13.9 | 97.0 |
| -3R+3K | 125 | 27250 | 13.9 | 97.0 |

## 3 Analysis of the simulation trajectories

### 3.1 Determining the size of a single chain

#### 3.1.1 Radius of gyration (R*_g_*)

To determine the size of the polymer chain, we computed R*_g_*, which is given by:

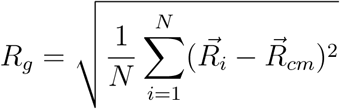

where *R_g_*is the radius of gyration, *R⃗_cm_* is the position vector of the center of mass, *R⃗_i_*is the position vector of the *i*^th^ bead, and *N* is the total number of beads.

#### 3.1.2 Hydrodynamic radius (R*_h_*)

The hydrodynamic radius of a specific protein conformation is defined by Nygaard et al., ^32^ who derived a sequence length (N) dependent relationship between the radius of gyration(R*_g_*) and R*_h_*

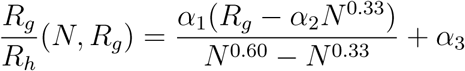

where, the fitting parameters are *α*_1_ = 0.216 Å*^−^*^1^, *α* = 4.06 Å, and *α*_3_ = 0.821.

### 3.2 Estimating Flory scaling exponent (*ν*)

The distance between *i^th^* and *j^th^*beads of a polymer chain can be denoted as:

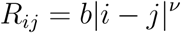

where *R_ij_*is the distance between the *i*^th^ and *j*^th^ beads, b is the Kuhn length(b=0.5).

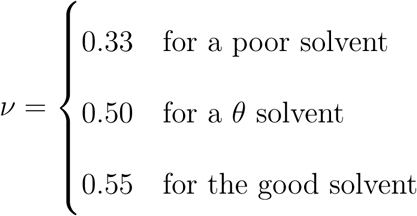

### 3.3 Estimation of scattering curves

We used the PULCHRA tool^33^ (a fast computational tool for converting coarse-grained protein models into all-atom protein structures) to convert coarse-grained simulation snapshots into atomistic representations. Then we used CRYSOL^34^ (a software program for evaluating the solution scattering from macromolecules with known atomic structure) to compute the SAXS intensity profiles. Traditionally, Kratky plots have been used to qualitatively assess the flexibility and/or degree of protein unfolding in samples. Unfolded (highly flexible) proteins exhibit a plateau in high q. In contrast, compact, globular proteins display a bell-shaped (Gaussian) peak. ^34,35^ A partially unfolded (flexible) protein may have a combination of the bell-shape and plateau, or a plateau that slowly decays to zero.

### 3.4 Estimation of critical point and the phase diagram

We simulate the slab box condensate over 1 *µ*s. Then, we compute the density profile along the Z-axis, as shown in Fig. 9. We determine the densities of the dense phase (*ρ_High_*) and the dilute phase(*ρ_Low_*) by fitting density profile as a sigmoidal function.

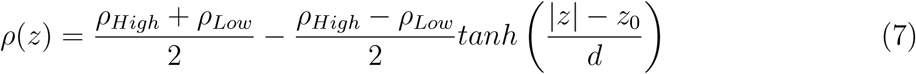

where *ρ*(*z*) is the density at position *z*, *z*_0_ is the phase boundary, and *d* is the thickness of the interface area. The critical point of the phase diagrams is determined by using the universal scaling law of coexistence densities near a critical point,^38^ and the law of rectilinear diameters.^39,40^

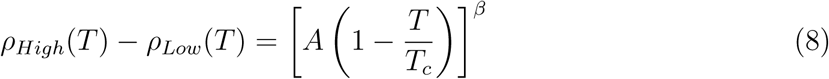

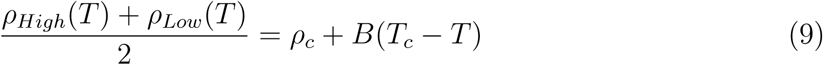

where *ρ_High_*(T) and *ρ_Low_*(T) are the densities of the dense and dilute phases of the condensate, respectively. The *T_c_* denotes the critical temperature and *ρ_c_* denotes the critical density extracted from fitting simulation data in equations 8 and 9. A and B are the fitting parameters. The parameter *β* is set to 0.5 according to the Flory-Huggins model for classical (mean-field) critical phenomena. ^41^

## 4 Results and Discussion

On simulating the four hnRNPA1-LCD variants (WT, Aro+, Aro−, and Aro−−) using the original SOP-IDP model,^20,21^ we find that the R*_g_* values of these four IDPs were approximately identical (*R_g_* ≈ 3.09 nm). In contrast, experimental R*_g_* values are noticeably sensitive to the aromatic valence patterning. This striking result illustrates that the SOP-IDP model is insensitive to the variations in aromatic residue content along the sequence. To make sure that the SOP-IDP model is truly insensitive to the aromatic residues content along the A1-LCD sequence, we estimated the *R_g_* values of some more A1-LCD variants (Fig. 1). The *R_g_* values were all in the range of 3.0 - 3.2 nm and insensitive to the aromatic valence patterning. When we attempted to form a biomolecular condensate using the slab box condensate simulation method, the original SOP-IDP model failed to do so. These observations indicate that the original SOP-IDP model cannot capture the influence of *π* −*π* and cation-*π* interactions at emergent scales.

### 4.1 Reparameterization of *π* − *π* and cation−*π* interactions

Cation−*π* interaction is stronger than *π* − *π* interactions^3^ and almost all IDP CG models honor this trend in their force field.^4,7,19,42^ In general, these models honor the following order of interaction strength *π* − *π*: *W W > Y Y > FF*, and the cation−*π* interaction of Arginine is more sticky than Lysine. However, when we examined the *π* − *π* and cation-*π* interactions in the original SOP-IDP model, which is derived from Betancourt-Thirumalai statistical potential,^26^ we found that the *π* − *π* interaction was stronger than the cation-*π* interaction (see Fig. 3(A) and (B). To improve the *π* −*π* and cation-*π* interactions, we follow the ordering of *π*−*π* and cation-*π* interaction according to the Mpipi forcefield. For the *π*−*π* interaction, the strength ordering was *FF > W F > W W > W Y > Y F > Y Y*, and for the cation−*π* interaction, the strength ordering was *RW > KY > RY > KW > RF > KF*. We also show the bar plot of *π* − *π* and cation-*π* interactions of SOP-Multi-*π* with the Mpipi force field in supporting Figure S1. To address this deficiency, we reparameterize the *π* − *π* and cation-*π* interactions within the SOP-IDP framework, yielding the revised model, hereafter referred to as SOP-Multi-*π*. A dimensionless rescaling prefactor *λ_ij_* was introduced as a multiplicative factor applied exclusively to aromatic *π* − *π* interaction pairs (FF, FW, FY, WY, WW, YY) and cation−*π* Interaction pairs (RF, RW, RY, KF, KW, KY). The optimal *λ_ij_* values (Fig. 3(C)) were determined through an iterative optimization procedure designed to minimize the root-mean-square error between simulated and experimental radii of gyration across A1-LCD and its 16 variants.

**Figure 3:**
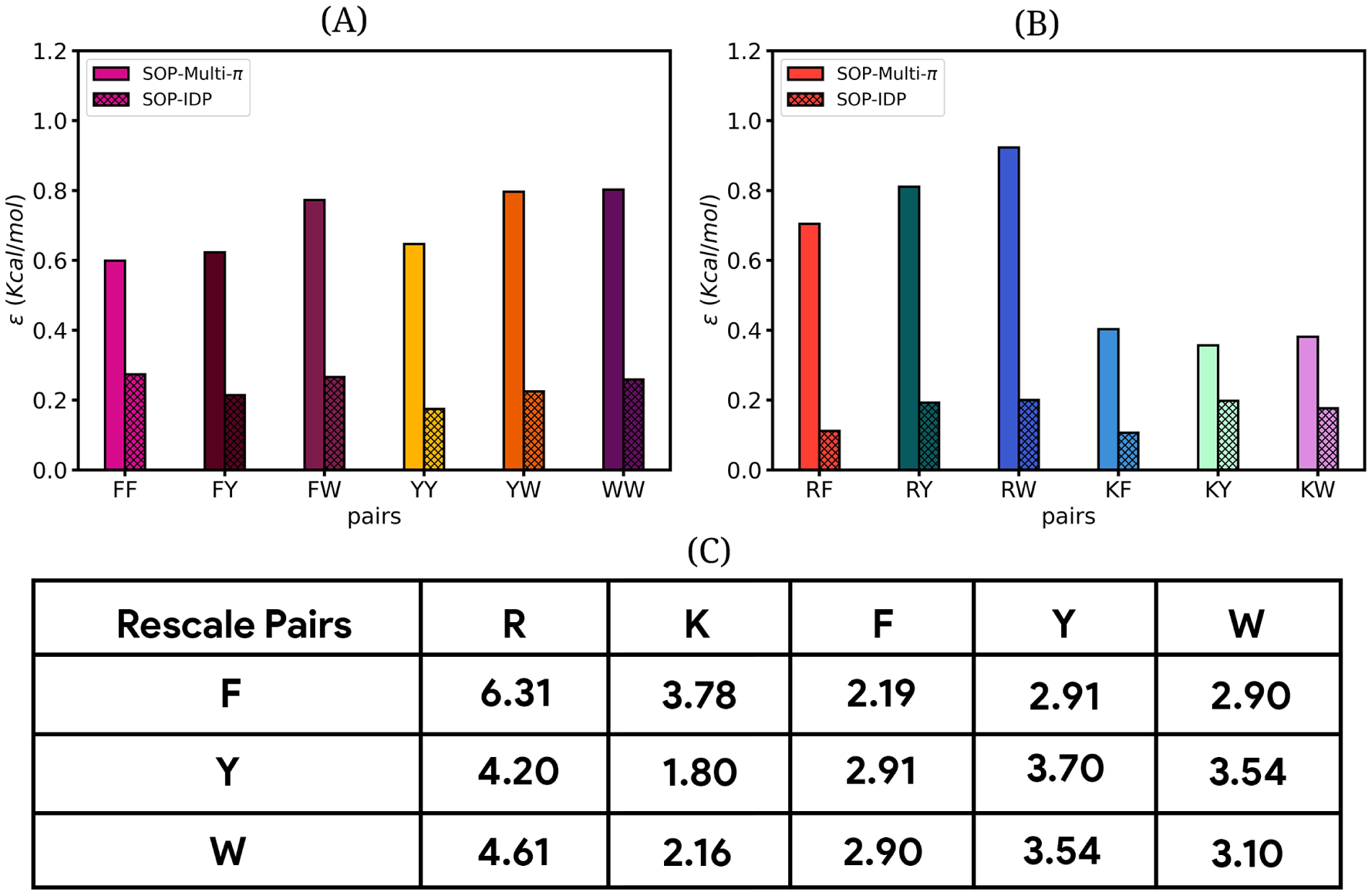
(A, B) The bar plot of pairwise interaction of *π* − *π* and cation−*π* pairs for SOP-IDP and SOP-Multi-*π* models. (C) Values of dimensionless rescale parameter *λ_ij_* for the different *π* − *π* and cation−*π* pairs

A physically meaningful constraint was imposed during reparameterization: Tyr–Tyr interactions were required to be stronger than Phe–Phe interactions. This hierarchy is consistent with the findings of De Sancho and López,^7^ and with other CG models. Notably, this ordering inverts the ranking observed in statistical potentials derived from folded-protein contact frequencies, in which Phe-Phe contacts predominate. This discrepancy is not paradoxical but rather reflects the fundamentally different chemical environment in solvent-exposed, disordered chains relative to the buried hydrophobic cores of folded proteins, a distinction that any physically appropriate IDP force field must account for. Following reparameterization, SOP-Multi-*π* yields radii of gyration of 2.65 ± 0.53, 2.54 ± 0.51, 2.86 ± 0.55, and 3.00 ± 0.57 nm for WT, Aro+, Aro−, and Aro−−, respectively, in excellent agreement with Guinier fits of experimental SEC-SAXS data. The probability density distributions of *R_g_* for all four variants are shown in Figure 4, where solid curves represent SOP-Multi-*π* results, dotted curves the SOP-IDP results, and vertical pink lines the experimental values. The marked differentiation across variants with varying aromatic content confirms that SOP-Multi-*π* has successfully recovered the sensitivity to aromatic identity that was absent in the SOP-IDP model. We estimated the radius of gyration, end-to-end distance, hydrodynamic radius, and Flory exponent of A1-LCD and its variants and report that in the supporting information Table S3. We compared the Flory exponent of A1-LCD and its variants with the SOP-IDP force field and found that SOP-IDP predicts IDP chains more extended than the SOP-Multi-*π* force field (Supporting Information Figure S2.)

**Figure 4:**
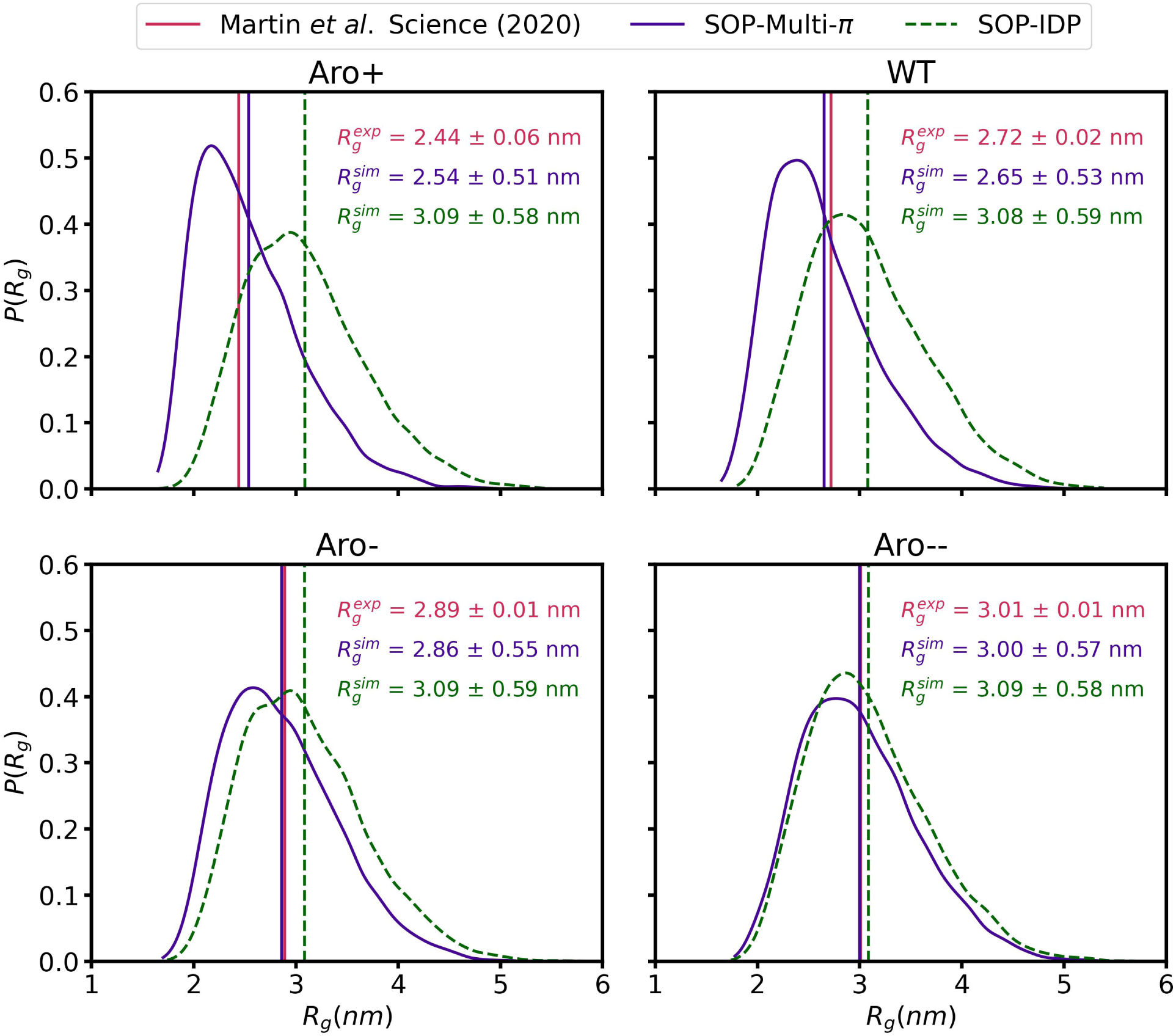
The probability density distribution of radii of gyration (*R_g_*) of WT, Aro+, Aro−, and Aro−−. The purple solid curves are SOP-Multi-*π* simulation results. The green dashed curves are SOP-IDP simulation results. The pink vertical line denotes the experimental *R_g_* values that were obtained from the Guinier fit of experimental SEC-SAXS data. The purple solid and dashed green vertical lines denote the mean *R_g_* values obtained with the SOP-Multi-*π* and SOP-IDP models, respectively. For convenience, the *R_g_* values are noted separately following the same color scheme.

We performed the Kratky analysis to assess whether our model accurately reproduces the conformational ensemble of the A1-LCD and its variants. Fig. 5 presents Kratky plots of A1-LCD and its variants, where the y-axis shows the dimensionless quantity (*qR_g_*)^2^ · *I*(*q*)*/I*_0_ and the x-axis shows *qR_g_*, a normalized representation that enables direct comparison of chain conformational properties across proteins of different sizes. Each panel contains three curves: the light grey noisy trace represents experimental SAXS (Small-Angle X-ray Scattering) data collected from solution, the green curve represents scattering profiles computed from coarse-grained molecular simulations using the SOP-IDP force field, and the pink curve represents profiles from the SOP-Multi-*π* force field. The 16 variants are systematically organized across four rows: the first row covers aromatic residue content variants (WT, Aro+, Aro−, and Aro−−), the second row covers Phenylalanine-to-Tyrosine swap variants (−12F+12Y, +7F−7Y, −9F+6Y, −8F+4Y), the third row mixes aromatic and Arg charge variants (−9F+3Y, +7R, +2R, −6R), and the fourth row systematically replaces Arginine with Lysine (−10R, −3R+3K, −6R+6K, −10R+10K) to isolate the contribution of cation-*π* interactions, since Arg carries a guanidinium group capable of *π*-stacking while Lys does not. Across most panels, SOP-IDP (green) tends to overestimate chain expansion, producing curves that rise steeply and remain elevated at high *qR_g_*, overshooting the experimental plateau. At the same time, SOP-Multi-*π* (pink) exhibits a more pronounced peak followed by a dip, capturing the transient compaction driven by *π* − *π* and cation−*π* contacts and agreeing more closely with the experimental data. Particularly in aromatic and Arg-rich variants with both models converging more closely in the Arg-depleted and Arg to Lys swap variants of Row 4, where *π*-mediated interactions are diminished, collectively demonstrating that explicitly accounting for *π* − *π* and cation−*π* interactions in SOP-Multi-*π* is critical for accurately reproducing the conformational ensemble of A1-LCD and its variants. We also compared the scattering intensity profiles from the simulation data with those from the experimental data. The scattering intensity profile has good agreement with the experimental data, as shown in the supporting information Figure S3.

**Figure 5:**
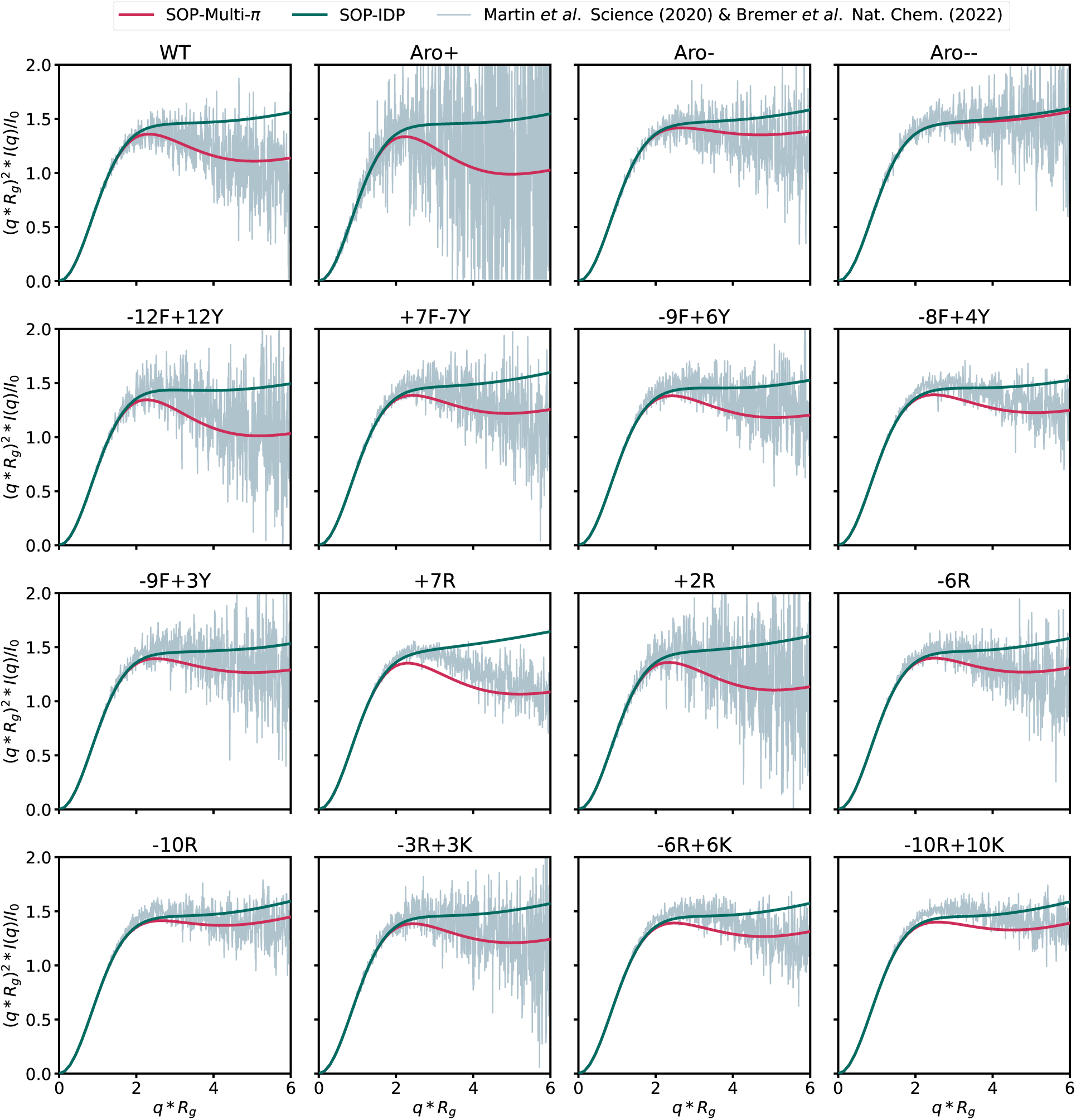
Kratky profile of A1-LCD and its varinats. The grey colour curve corresponds to the experimental data, green colour denotes the scattering curve obtained from the SOP-IDP simulation, and the pink colour curve denotes the scattering profile obtained from the SOP-Multi-*π* simulation.

### 4.2 SOP-Multi-*π* exhibits robust behavior across a range of IDPs

We assessed the quantitative performance for R*_g_* and SAXS profile between simulated and experimental values for all the above-mentioned systems (all sequences in Fig. 1). For the original SOP-IDP model, the Pearson correlation for R*_g_* is quite poor at r = 0.07, with a root-mean-square deviation (D) = 0.40 nm (Figure 6a). SOP-Multi-*π* force field shows excellent improvement in correlation between simulated and experimental radii of gyration. We get Pearson’s correlation (r) = 0.83 and root-mean-square deviation (D) = 0.08 nm, shown in (Figure 6b), a substantial improvement. To validate the robustness of the reparameterized interaction, we have simulated a diverse set of 34 other IDPs spanning lengths from 24 to 198 residues and calculated the R*_g_* values from the simulations. The reparameterization generalizes well beyond the training set without overfitting, yielding r = 0.93 (Figure c). Flory scaling analysis (Figure 6d) further demonstrates that simulated chains follow the expected *ν* ≈ 0.588 exponent characteristic of good-solvent behavior, consistent with the experimentally characterized expanded conformations of IDPs. We report the radius of gyration, end-to-end distance, hydrodynamic radius, and Flory exponent of 34 IDPs in the supporting information Table S4.

**Figure 6:**
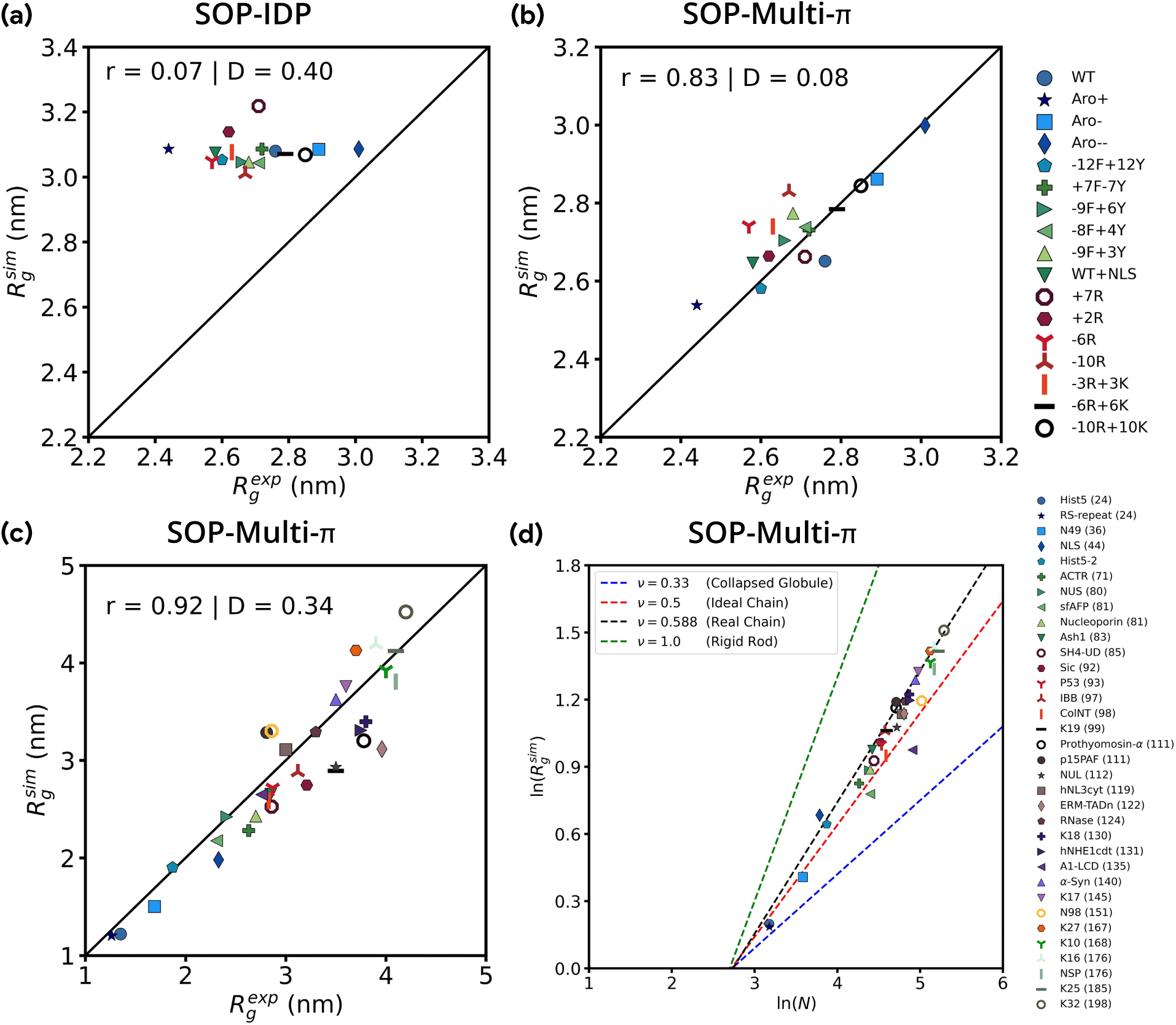
The Pearson’s correlation coefficient (r) and root mean squared deviation (D) between experimental *R_g_* values and those estimated from simulation for A1-LCD (WT) and its 16 variants, for (a) SOP-IDP model, (b) SOP-Multi-*π* model. (c) Pearson’s correlation between simulated and experimental *R_g_* values for a dataset consisting of 34 IDPs with chain length varying from 24 to 198 residues. These simulations were carried out at given experimental temperature and salt concentration conditions using the SOP-Multi-*π* force field. In (a)-(c), the black diagonal line serves as a guide to the eye and denotes a perfect match with the experiment. (d) The *R_g_* values obtained from SOP-Multi-*π* simulations fitted to the Flory scaling law *R_g_* ∝ *N ^ν^* (shown as dashed lines). The exponent *ν* = 0.588, consistent with polymer in a good solvent behavior observed for many IDPs.

SOP-Multi-*π* represents a dramatic and meaningful improvement over SOP-IDP force field, as demonstrated by the contrast between panels 6(a) and 6(b). Panel 6(c) extends the benchmark to a diverse set of 34 different IDPs, ranging from short Histatin-5 IDP (24 residues) to large disordered chains (198 residues) with impressive correlation at r = 0.93. Panel 6(d) presents the ln(*R_g_*) versus ln(N) scaling plot, where data from the diverse IDP set aligns well with the dashed line corresponding to a Flory exponent of *ν* = 0.588, the value expected for a self-avoiding random walk in a good solvent. Taken together, SOP-Multi-*π* is a substantially improved and physically more grounded model that successfully captures sequence-sensitive compaction across a broad chemical and chain-length space. Still, the persistent absolute errors, systematic short-peptide overestimation, and residual outliers at extreme sequence compositions collectively indicate that the force field is not yet fully transferable and that further refinement, particularly in how it handles short disordered peptides and sequences with highly atypical charge or aromatic distributions, will be necessary before it can be applied with full confidence across the complete IDP landscape.

### 4.3 SOP-Multi-*π* captures the single-Chain properties of multi-domain proteins

SOP-Multi-*π* captures the global conformational properties and experimentally observed scattering characteristics of several multidomain, intrinsically disordered RNA-binding proteins, including hnRNPA1, TDP-43, G3BP1, and FUS. The domain architectures are illustrated in Fig. 7A. As is evident from Fig. 7 B, C the mean simulated radius of gyration 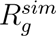 = 3.29 ± 0.66 and 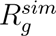 = 3.86 ± 0.79 nm of hnRNPA1 and TDP43 agrees well with the experimental value 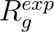 = 3.12 and 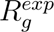 = 4.19 nm, respectively. The model can also capture the salt-dependent conformational behavior of G3BP1. Fig. 8 compares the simulation-derived SAXS profiles of the full-length hnRNPA1, TDP43, G3BP1, and FUS multidomain proteins against the experimental SAXS data. SAXS comparison data clearly demonstrate that these conformational ensembles generated by SOP-Multi-*π* force fields yield realistic small-angle X-ray scattering (SAXS) profiles. For hnRNPA1 and TDP-43, the simulated scattering curves reproduce the overall experimental intensity decay and characteristic features in both the conventional and Kratky representations, while simulations of G3BP1 recapitulate the experimentally observed changes in scattering behavior upon increasing salt concentration. The FUS simulations similarly yield a physically consistent Kratky profile, despite its highly disordered, multidomain architecture. Overall, the agreement between both *R_g_* distributions and SAXS profiles indicates that the coarse-grained framework can capture the essential dimensions, conformational heterogeneity, and large-scale structural organization of complex multidomain proteins, supporting its applicability to describing experimentally relevant ensembles of intrinsically disordered and phase-separating proteins.

**Figure 7:**
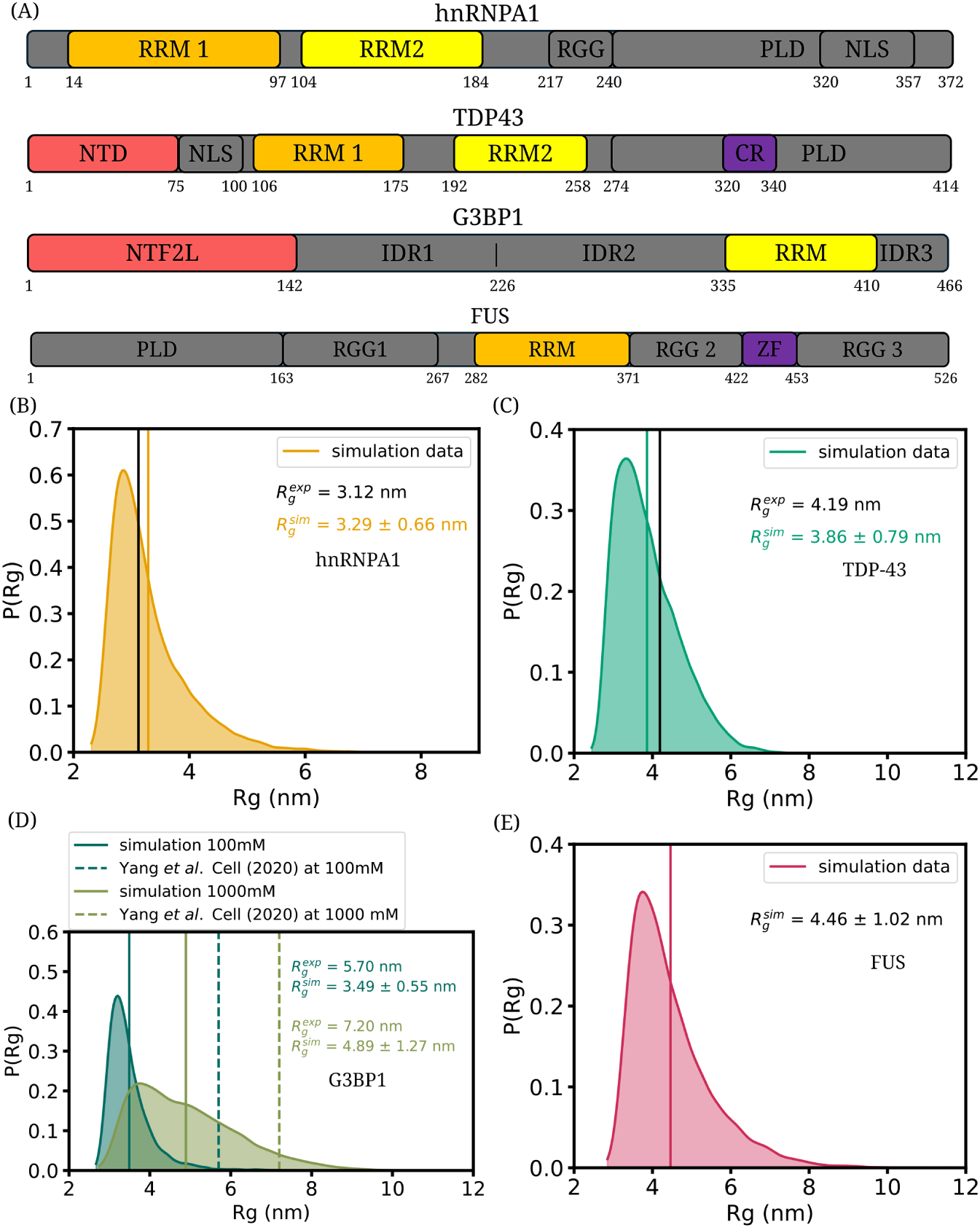
(A)Different sequence domains of four proteins hnRNPA1, TDP43, G3BP1, and FUS: RNA recognition motifs (RRM), arginine-glycine-rich regions (RGG), prion-like domain (PLD), nuclear localization sequence (NLS), N-tail domain (NTD), *α*-helical region denoted as conserved region (CR), nuclear transport factor 2-like (NTF2L) domain, Intrinsically disordered region (IDR), and zinc finger (ZF). (B) and (C) Probability distributions of the radius of gyration (*R_g_*) for hnRNPA1 and TDP-43, respectively. The orange and green vertical lines indicate the mean simulated 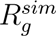 values for hnRNPA1 and TDP-43, respectively, while the solid black vertical line represents the experimental 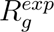. (D) Probability distribution of *R_g_* for G3BP1 at 100 and 1000mM salt concentration in green and olive colour, respectively. The solid and dashed vertical lines at 100 and 1000 mM salt concentrations, corresponding to the green and olive colours, denote 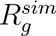 and 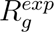, respectively. (E) Probability distribution of R*_g_* for FUS, and the vertical pink line denotes the mean 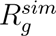 value of simulated data.

**Figure 8:**
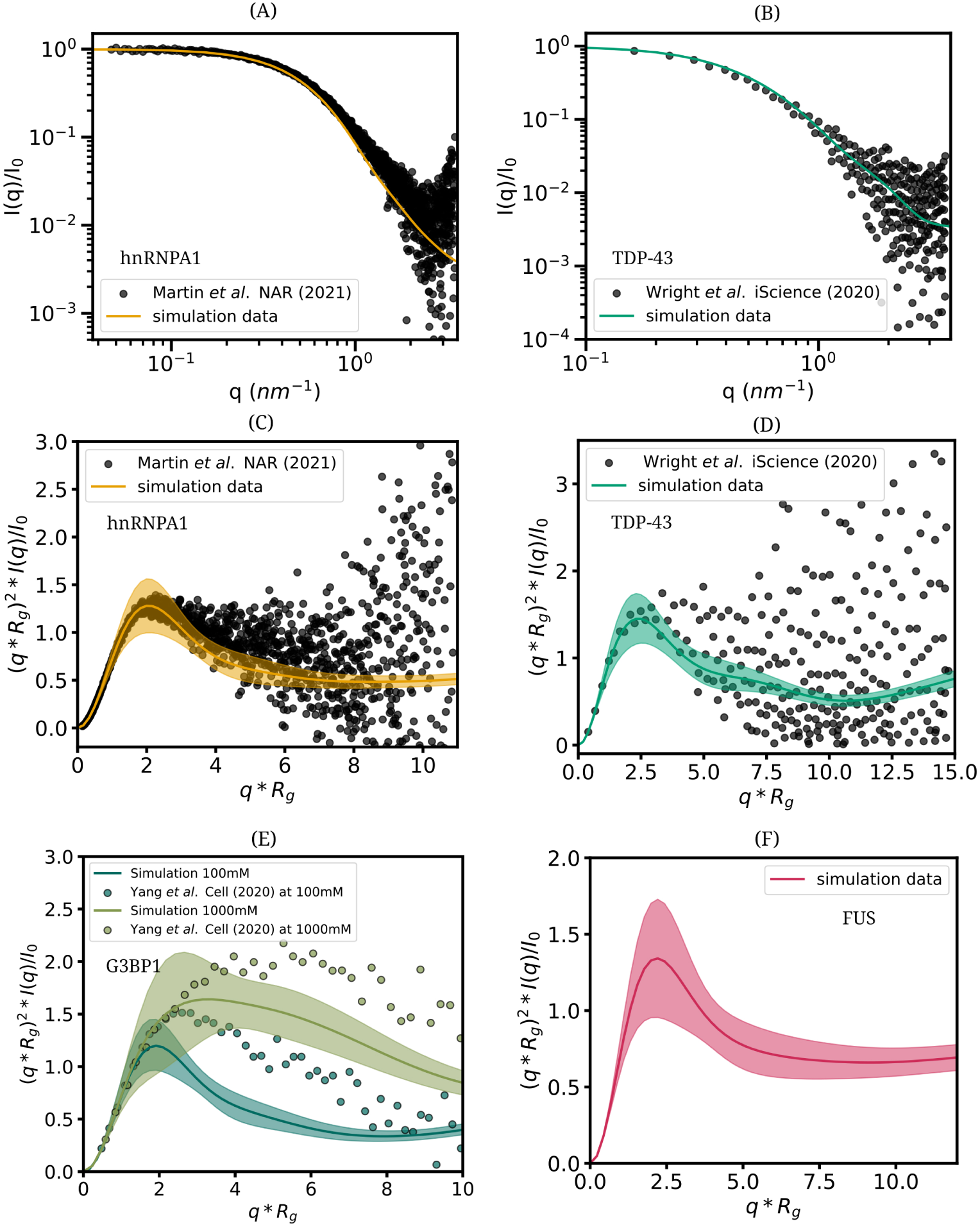
(A) and (B) SAXS curves of hnRNPA1 and TDP-43 were computed from simulation data (orange and green curves, respectively) and overlaid on the experimental data (black dots). (C) and (D) SAXS curve in (A) and (B) shown in the Kratky representation for hnRNPA1 and TDP-43, respectively. (E) The SAXS curve in the Kratky representation was computed from simulation data for the G3BP1 protein at two salt concentrations (100 and 1000 mM), shown in green and olive, respectively, and overlaid on the experimental data (corresponding dots). (F) The SAXS curve shown in the Kratky representation was computed from simulation data for the FUS protein.

### 4.4 SOP-Multi-*π* accurately captures the phase behavior of IDPs

It is quite challenging for a highly coarse-grained force field to recapitulate both the dilute solvent phase single-chain behavior and condensed phase behavior. This problem is particularly exacerbated in sequences with a substantial fraction of aromatic residues, which can engage in *π*-*π* and cation-*π* contacts. For instance, the HPS-cation-*π* force field, which uses an auxiliary term for describing aromatic interaction, severely underestimates the sizes of IDPs, such as *α*-synuclein, while providing a reasonably accurate description of its phase behavior. It is thus natural to ask if the SOP-Multi-*π* model, primarily optimized with respect to single-chain properties, could probe phase separation. This is especially important since the original SOP-IPD model exhibits no phase separation behavior at all. As we describe below, the SOP-Multi-*π* model provides a fairly accurate description of the phase behavior of A1-LCD and its variants. Our findings illustrate how a simple rescaling of *π*-*π* and cation-*π* interactions, derived from a statistical potential rather than a full-fledged reparametrization, could be sufficient to restore the balance in an otherwise well-behaved force field.

In Fig 9, we show the slab density profile at different temperatures obtained from the direct coexistence simulation. We obtained a flat-density profile using the SOP-IDP force field for A1-LCD and its three variants. In contrast, a dense phase forms at the center of the slab box when using the SOP-Multi-*π* force field. As expected, the density profile flattens and mixing occurs as the temperature increases.

**Figure 9:**
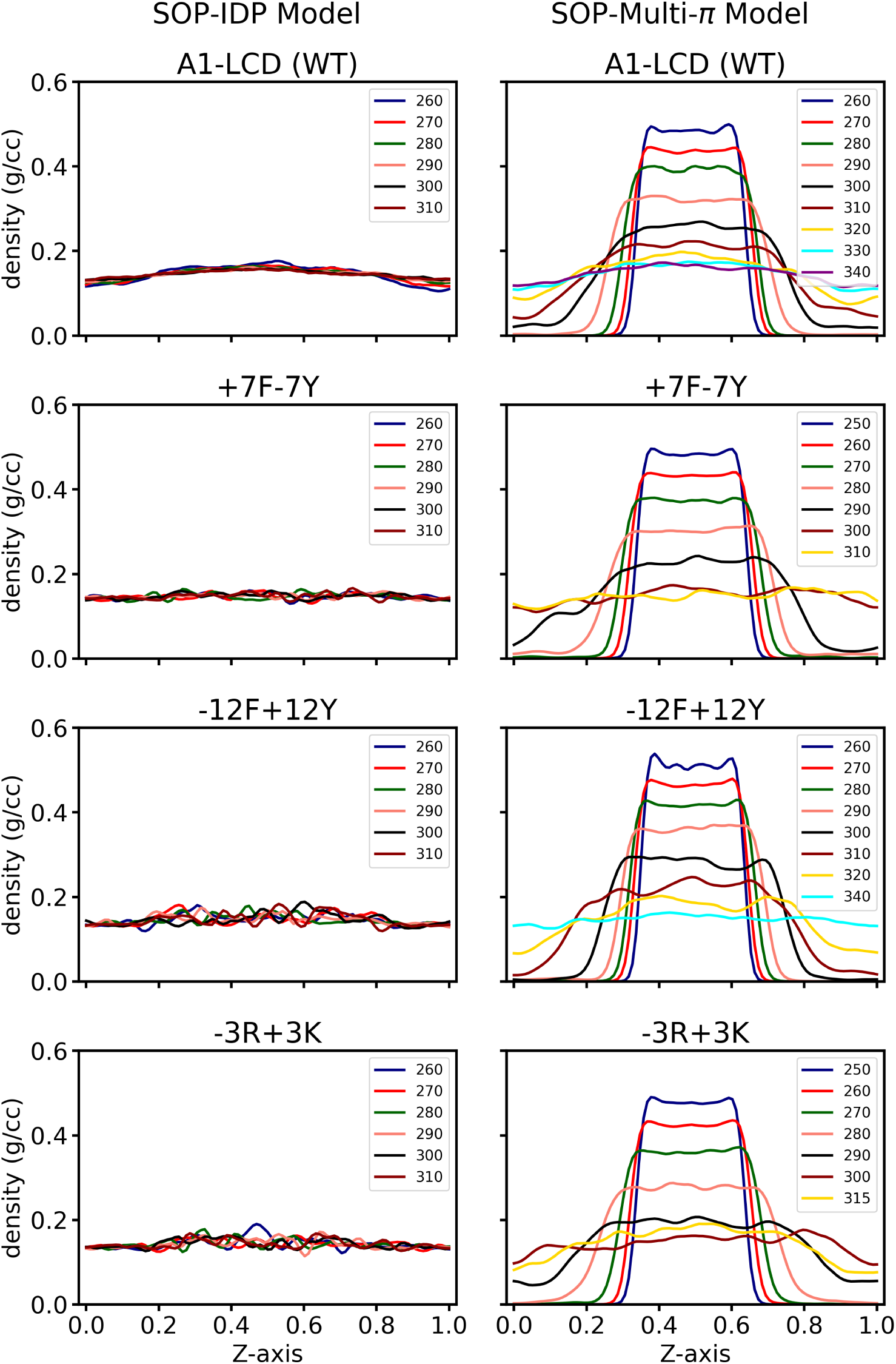
The slab density profile from direct coexistence simulation for A1-LCD and its three variants (−12F+12Y, +7F−7Y, and −3R+3K). Result obtained with SOP-IDP shown on the left panel and with SOP-Multi-*π* on the right panel.

The phase diagram of A1-LCD and some of its variants is shown in Fig. 10. The width of the two-phase regime is greater for the −12F+12Y variant than for A1-LCD(WT), as shown in Fig. 10(B). Tyrosine has more stickers than Phenylalanine, so when all phe (F) was replaced by Tyr (Y), the two-phase regime of −12F+12Y is widened, and the right arm of the binodal curve is shifted to higher concentrations than in A1-LCD. This widening indicates that replacing Phe(F) residues with Tyr(Y) residues enhances the driving force for phase separation. The width of the two-phase regime was narrowed for the +7F−7Y variant (all Tyr (Y) was replaced by Phe (F)) compared to A1-LCD. The right arm of the binodal curve for +7F−7Y is shifted to a lower concentration than A1-LCD. The −3R+3K variant has 3 Arg (R) residues, which were replaced by Lys (K) residues. The two-phase regime of −3R+3K is narrower than that of A1-LCD. The right-hand arm (dense phase region) of the binodal curve shifted towards the lower concentration because Lys (K) is a weaker sticker than Arg (R).^3^ We conclude that only reparameterizing the *π*−*π* and cation-*π* interactions at the single-molecule level in the SOP-IDP model allows us to capture the molecular assembly behavior very well.

**Figure 10:**
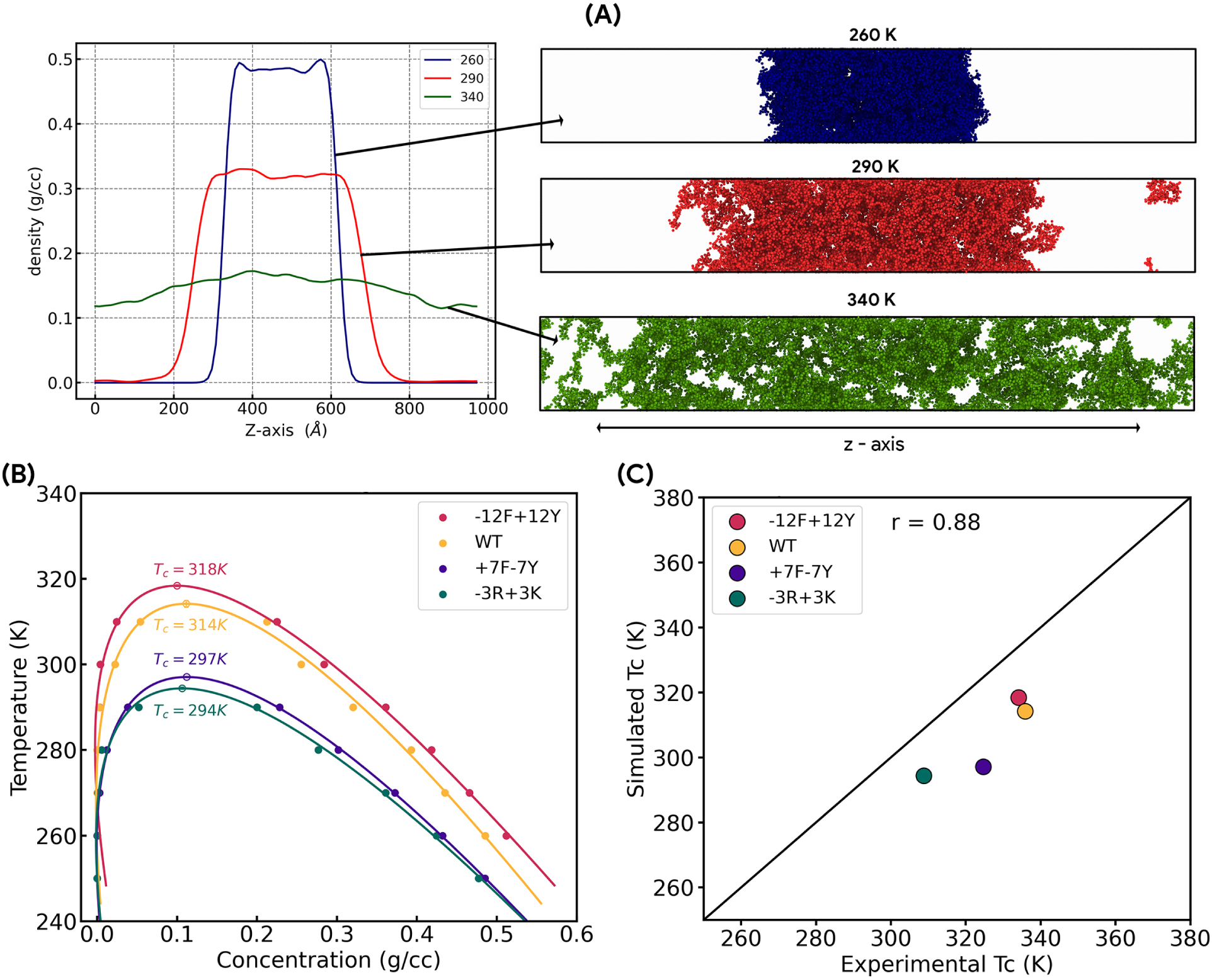
(A) Slab density profile at a set of temperatures and configurations of the A1-LCD slab simulation is shown on the right-hand side. (B) Phase diagram of the A1-LCD and its three variants (−12F+12Y, +7F−7Y, and −3R+3K). The Flory-Huggins theory fitted critical temperatures of −12F+12Y, WT, +7F−7Y, and −3R+3K are 318K, 314K, 297K, and 294K, respectively. (C) A comparison of the critical temperature (T*_c_*) determined by simulation data and the theoretically predicted experimental critical temperature (T*_c_*).

The SOP-Multi-*π* model captures both the characterization of single-chain molecule IDPs and the emergent behavior of IDPs. Despite the noticeable improvement, the SOP-Multi-*π* force field somewhat underestimates the experimentally predicted critical temperature, suggesting that the treatment of *π* − *π* and cation-*π* interactions is not yet fully optimal within the limits of CGFF parametrization.

### 4.5 Discussion

In this work, we present SOP-Multi-*π*. This enhanced two-bead-per-amino-acid coarse-grained model systematically addresses the central deficiency of the original SOP-IDP framework: its insensitivity to *π* − *π* and Cation-*π* interactions. The primary finding is that the single-molecule conformational properties and condensate phase behavior of aromatic-rich IDPs are governed by a relatively small but physically critical set of residue-pair interactions specifically those involving Phe, Tyr, Trp, Arg, and Lys. We show that accurately weighting these interactions via dimensionless rescaling prefactors *λ_ij_* is sufficient to achieve quantitative agreement with experiments across a broad sequence and length space. A key conceptual result is that condensate formation in A1-LCD requires no parameters beyond those already present in SOP-IDP, only the correct weighting of the *π* −*π* and cation-*π* terms. This is consistent with the molecular grammar framework proposed by Wang et al., ^3^ wherein aromatic stickers and disordered spacers jointly determine phase behavior. The reparameterization confirms the primacy of Tyr–Tyr interactions in driving condensate formation, placing Tyr above Phe in the interaction hierarchy in agreement with single-chain and condensate simulations by De Sancho and López,^7^ and with the Mpipi,^17^ CALVADOS,^42^ and HPS-Urry^19^ models. This ranking stands in explicit contrast to statistical potentials derived from folded protein contacts, in which Phe–Phe contacts predominate. This discrepancy reflects the fundamentally different chemical environment of solvent-exposed disordered chains compared with folded protein interiors.

The sequential parameterization strategy, calibrating single-chain properties first, then transferring the same parameters to condensate simulations, provides an important and non-trivial validation. The direct transfer of force-field parameters from single-chain *R_g_* optimization (A1-LCD and its 16 variants) to condensate critical temperature prediction (r = 0.88) implies a deep physical consistency: the residue-level interactions governing chain compaction are the same interactions stabilizing the dense condensate phase. This cross-scale transferability is an encouraging and non-obvious property of the rescaled model, supporting its physical rather than purely empirical basis. Validation across 34 IDPs (r = 0.93 on *R_g_*) and across multidomain proteins containing folded domains and IDRs, notably hnRNPA1, FUS, G3BP1 and TDP-43, establishes that SOP-Multi-*π* is not overfit to the A1-LCD training set. The ability to back-map CG ensembles to all-atom representations via PULCHRA^33^ and compute SAXS profiles with CRYSOL^34^ provides a direct experimental link that is unavailable in most single-bead CG models, making SOP-Multi-*π* particularly well-suited for comparison with SAXS, NMR, and single-molecule datasets.

### Scope and Limitation

A critical appraisal of the model’s limitations is warranted. The most pronounced discrepancy was observed for G3BP1, where trends related to salt-dependent changes in R*_g_* was observed but R*_g_* values were underestimated by ≈ 2.21-2.31 nm at 100 and 1000mM NaCl. G3BP1 is physiologically dimeric, and its unusually extended IDR conformation is likely driven by electrostatic repulsion and by NTF2L dimerization geometry, effects not captured in monomer simulations that employ a Debye-Hückel electrostatic treatment. More broadly, implicit-solvent models may systematically underestimate electrostatically driven chain expansion at low ionic strength, a limitation that future work should address through explicit coarse-grained water representations or improved dielectric screening functions. The quantitative accuracy observed for A1-LCD variants (r = 0.88) arises in part because relative comparisons within a shared sequence background benefit from partial error cancellation. Extending quantitative absolute critical-temperature predictions to diverse protein sequences will likely require temperature-dependent parameterization or an improved treatment of hydrophobic effects with advanced non-polar solvation features in the model.

Despite the limitations, SOP-Multi-*π* occupies a well-defined niche in the current landscape of IDP and condensate simulation tools. Its two-bead resolution enables microsecond-scale single-chain and condensate simulations, while PULCHRA back-mapping provides all-atom SAXS profiles for direct experimental validation, a combination unavailable in lower-resolution single-bead models. The model is computationally efficient, fully implemented in LAMMPS.^29^ We provide a user manual for SOP-Multi-*π* as a separate supporting information file. We believe that SOP-Multi-*π* will serve as a broadly applicable tool for investigating the sequence determinants of IDP conformational behavior and biomolecular condensate formation, as well as for generating testable structural predictions for experiments.

## Acknowledgement

AS acknowledges financial support from the Indian Institute of Science (IISc) and the high-performance computing facility “Beagle,” which was set up with grants from the erstwhile IISc-DBT partnership program. SP thanks the Ministry of Education, Government of India, for the graduate fellowship. AS acknowledges the FIST program sponsored by the Department of Science and Technology, India, that supports the MBU infrastructure. AS thanks the DBT National Network Project (NNP) grant (BT/PR40323/BTIS/137/78/2023), the Matrics grants (MTR/2023/001040) and the ARG grant (ANRF/ARG/2025/009141/LS) from ANRF, India. AS and TJP would also like to thank the Teams Science Grant from the DBT-Wellcome Trust India Alliance (Grant number: IA/TSG/21/1/600245). DC acknowledges funding from ANRF through the Research grant (ANRF/ECRG/2024/003762/CS). SP thanks the Ministry of Education, Government of India, for the graduate fellowship.

## Author contributions

AS conceived the project and designed the research with SP. SP developed the rescaled SOP-Multi-*π* force field under the supervision of AS and DC. SP carried out single-chain and condensate IDPs simulations and analyzed their physical properties. NM verified simulations on multi-domain proteins and prepared the comprehensive manual that can be used to create LAMMPS data files for condensates with multiple types of full-length proteins (Supporting Information S2). AS and DC supervised the study. SP prepared the first draft of the paper. NM, DC, and AS polished the draft.

## Data availability statement

Our code, models, and curated datasets are publicly available at the Zenodo repository

## Supporting Information (Additional Data)

### The sequence of hnRNPA1-LCD and its mutants

We selected 16 mutant systems of hnRNPA1-LCD from the experimental work of Tanja Mittag and co-workers^1,2^ (residues 186-320, (UniProt: P09651); Isoform A1-A). All 16 sequences are listed below.

#### A-LCD (WT)

MASASSSQRG RSGSGNFGGG RGGGFGGNDN FGRGGNFSGR GGFGGSRGGG GYGGSGDGYN GFGNDGSNFG GGGSYNDFGN YNNQSSNFGP MKGGNFGGRS SGGSGGGGQY FAKPRNQGGY GGSSSSSSYG SGRRF

#### A-LCD^+NLS^

MASASSSQRG RSGSGNFGGG RGGGFGGNDN FGRGGNFSGR GGFGGSRGGG GYGGSGDGYN GFGNDGSNFG GGGSYNDFGN YNNQSSNFGP MKGGNFGGRS SGPYGGGGQY FAKPRNQGGY GGSSSSSSYG SGRRF

#### Aro+

MAFASSFQRG RYGSGNFGGG RGGGFGGNDN FGRGGNFSGR GGFGGSRGGG GYGGSGDGYN GFGNDGSNFG GGGSYNDFGN YNNQSSNFGP MKGGNFGGRS SGGSYGGGQY FAKPRNQGGY GGSSFSSSYG SGRRF

#### Aro−

MASASSSQRG RSGSGNSGGG RGGGFGGNDN FGRGGNSSGR GGFGGSRGGG GYGGSGDGYN GFGNDGSNSG GGGSSNDFGN YNNQSSNFGP MKGGNFGGRS SGGSGGGGQY SAKPRNQGGY GGSSSSSSSG SGRRF

#### Aro−−

MASASSSQRG RSGSGNSGGG RGGGFGGNDN SGRGGNSSGR GGFGGSRGGG GSGGSGDGYN GSGNDGSNSG GGGSSNDFGN SNNQSSNSGP MKGGNFGGRS SGGSGGGGQY SAKPRNQGGS GGSSSSSSSG SGRRS

#### +7F−7Y

MASASSSQRG RSGSGNFGGG RGGGFGGNDN FGRGGNFSGR GGFGGSRGGG GFGGSGDGFN GFGNDGSNFG GGGSFNDFGN FNNQSSNFGP MKGGNFGGRS SGGSGGGGQF FAKPRNQGGF GGSSSSSSFG SGRRF

#### −12F+12Y

MASASSSQRG RSGSGNYGGG RGGGYGGNDN YGRGGNYSGR GGYGGSRGGG GYGGSGDGYN GYGNDGSNYG GGGSYNDYGN YNNQSSNYGP MKGGNYGGRS SGGSGGGGQY YAKPRNQGGY GGSSSSSSYG SGRRY

#### −9F+6Y

MASASSSQRGR SGSGNFGGG RGGGYGGNDN YGRGGNYSGR GGFGGSRGGG GYGGSGDGYN GGGNDGSNYG GGGSYNDSGN YNNQSSNFGP MKGGNYGGRS SGGSGGGGQY GAKPRNQGGY GGSSSSSSYG SGRRY

#### −8F+4Y

MASASSSQRGR SGSGNFGGG RGGGYGGNDN GGRGGNYSGR GGFGGSRGGG GYGGSGDGYN GGGNDGSNYG GGGSYNDSGN YNNQSSNFGP MKGGNYGGRS SGGSGGGGQY GAKPRNQGGY GGSSSSSSYG SGRRF

#### −9F+3Y

MASASSSQRG RSGSGNFGGG RGGGYGGNDN GGRGGNYSGR GGFGGSRGGG GYGGSGDGYN GGGNDGSNYG GGGSYNDSGN GNNQSSNFGP MKGGNYGGRS SGGSGGGGQY GAKPRNQGGY GGSSSSSSYG SGRRS

#### +7R

MASASSSQRG RSGRGNFGGG RGGGFGGNDN FGRGGNFSGR GGFGGSRGGG RYGGSGDRYN GFGNDGRNFG GGGSYNDFGN YNNQSSNFGP MKGGNFRGRS SGPYGRGGQY FAKPRNQGGY GGSSSSRSYG SGRRF

#### +2R

MASASSSQRG RSGSGNFGGG RGGGFGGNDN FGRGGNFSGR GGFGGSRGGG GYGGSGDGYN GFRNDGSNFG GGGRYNDFGN YNNQSSNFGP MKGGNFGGRS SGPYGGGGQY FAKPRNQGGY GGSSSSSSYG SGRRF

#### −6R

MASASSSQGG RSGSGNFGGG RGGGFGGNDN FGGGGNFSGS GGFGGSRGGG GYGGSGDGYN GFGNDGSNFG GGGSYNDFGN YNNQSSNFGP MKGGNFGGSS SGPYGGGGQY FAKPGNQGGY GGSSSSSSYG SGGRF

#### −10R

MASASSSQGG SSGSGNFGGG GGGGFGGNDN FGGGGNFSGS GGFGGSGGGG GYGGSGDGYN GFGNDGSNFG GGGSYNDFGN YNNQSSNFGP MKGGNFGGSS SGPYGGGGQY FAKPGNQGGY GGSSSSSSYG SGGGF

#### −3R+3K

MASASSSQRG KSGSGNFGGG RGGGFGGNDN FGRGGNFSGR GGFGGSKGGG GYGGSGDGYN GFGNDGSNFG GGGSYNDFGN YNNQSSNFGP MKGGNFGGRS SGGSGGGGQY FAKPRNQGGY GGSSSSSSYG SGRKF

#### −6R+6K

MASASSSQKG KSGSGNFGGG RGGGFGGNDN FGKGGNFSGR GGFGGSKGGG GYGGSGDGYN GFGNDGSNFG GGGSYNDFGN YNNQSSNFGP MKGGNFGGKS SGGSGGGGQY FAKPRNQGGY GGSSSSSSYG SGRKF

#### −10R+10K

MASASSSQKG KSGSGNFGGG KGGGFGGNDN FGKGGNFSGK GGFGGSKGGG GYGGSGDGYN GFGNDGSNFG GGGSYNDFGN YNNQSSNFGP MKGGNFGGKS SGGSGGGGQY FAKPKNQGGY GGSSSSSSYG SGKKF

### The sequence of 34 different lengths of IDPs

#### Histatin5

DSHAKRHHGY KRKFHEKHHS HRGY

#### RS-repeat

GAMGPSYGRS RSRSRSRSRS RSRS

#### N49

GCQTSRGLFG NNNTNNINNS SSGMNNASAG LFGSKP

#### NLS

ACETNKRKRE QISTDNEAKM QIQEEKSPKK KRKKRSSKAN KPPE

#### Histatin5-2

DSHAKRHHGY KRKFHEKHHS HRGYDSHAKR HHGYKRKFHE KHHSHRGY

#### ACTR

GTQNRPLLRN SLDDLVGPPS NLEGQSDERA LLDQLHTLLS NTDATGLEEI DRALGIPELV NQGQALEPKQ D

#### NUS

GCPSASPAFG ANQTPTFGQS QGASQPNPPG FGSISSSTAL FPTGSQPAPP TFGTVSSSSQ PPVFGQQPSQ SAFGSGTTPN

#### sfAFP

CKGADGAHGV NGCPGTAGAA GSVGGPGCDG GHGGNGGNGN PGCAGGVGGA GGASGGTGVG GRGGKGGSGT PKGADGAPGA P

#### Nucleoporin

GCPSASPAFG ANQTPTFGQS QGASQPNPPG FGSISSSTAL FPTGSQPAPP TFGTVSSSSQ PPVFGQQPSQ SAFGSGTTPN A

#### Ash1

GASASSSPSP STPTKSGKMR SRSSSPVRPK AYTPSPRSPN YHRFALDSPP QSPRRSSNSS ITKKGSRRSS GSSPTRHTTR VCV

#### SH4-UD

MGSNKSKPKD ASQRRRSLEP AENVHGAGGG AFPASQTPSK PASADGHRGP SAAFAPAAAE PKLFGGFNSS DTVTSPQRAG PLAGG

#### Sic1

MTPSTPPRSR GTRYLAQPSG NTSSSALMQG QKTPQKPSQN LVPVTPSTTK SFKNAPLLAP PNSNMGMTSP FNGLTSPQRS PFPKSSVKRT

#### P53

MEEPQSDPSV EPPLSQETFS DLWKLLPENN VLSPLPSQAM DDLMLSPDDI EQWFTEDPGP DEAPRMPEAA PPVAPAPAAP TPAAPAPAPS WPL

#### IBB

GCTNENANTP AARLHRFKNK GKDSTEMRRR RIEVNVELRK AKKDDQMLKR RNVSSFPDDA TSPLQENRNN QGTVNWSVDD IVKGINSSNV ENQLQAT

#### ColNT

MGSNGADNAH NNAFGGGKNP GIGNTSGAGS NGSASSNRGN SNGWSWSNKP HKNDGFHSDG SYHITFHGDN NSKPKPGGNS GNRGNNGDGA SSHHHHHH

#### K19

MQTAPVPMPD LKNVKSKIGS TENLKHQPGG GKVQIVYKPV DLSKVTSKCG SLGNIHHKPG GGQVEVKSEK LDFKDRVQSK IGSLDNITHV PGGGNKKIE

#### Prothymosin-*α*

MSDAAVDTSS EITTKDLKEK KEVVEEAENG RDAPANGNAE NEENGEQEAD NEVDEEEEEG GEEEEEEEEG DGEEEDGDED EEAESATGKR AAEDDEDDDV DTKKQKTDED D

#### p15PAF

MVRTKADSVP GTYRKVVAAR APRKVLGSST SATNSTSVSS RKAENKYAGG NPVCVRPTPK WQKGIGEFFR LSPKDSEKEN QIPEEAGSSG LGKAKRKACP LQPDHTNDEK E

#### NUL

GCGFKGFDTS SSSSNSAASS SFKFGVSSSS SGPSQTLTST GNFKFGDQGG FKIGVSSDSG SINPMSEGFK FSKPIGDFKF GVSSESKPEE VKKDSKNDNF KFGLSSGLSN PV

#### hNL3cyt

MYRKDKRRQE PLRQPSPQRG AWAPELGAAP EEELAALQLG PTHHECEAGP PHDTLRLTAL PDYTLTLRRS PDDIPLMTPN TITMIPNSLV GLQTLHPYNT FAAGFNSTGL PHSHSTTRV

#### ERM-TADn

MDGFYDQQVP FMVPGKSRSE ECRGRPVIDR KRKFLDTDLA HDSEELFQDL SQLQEAWLAE AQVPDDEQFV PDFQSDNLVL HAPPPTKIKR ELHSPSSELS SCSHEQALGA NYGEKCLYNY CA

#### RNaseA

KETAAAKFER QHMDSSTSAA SSSNYCNQMM KSRNLTKDRC KPVNTFVHES LADVQAVCSQ KNVACKNGQT NCYQSYSTMS ITDCRETGSS KYPNCAYKTT QANKHIIVAC EGNPYVPVHF DASV

#### K18

MQTAPVPMPD LKNVKSKIGS TENLKHQPGG GKVQIINKKL DLSNVQSKCG SKDNIKHVPG GGSVQIVYKP VDLSKVTSKC GSLGNIHHKP GGGQVEVKSE KLDFKDRVQS KIGSLDNITH VPGGGNKKIE

#### hNHE1cdt

MVPAHKLDSP TMSRARIGSD PLAYEPKEDL PVITIDPASP QSPESVDLVN EELKGKVLGL SRDPAKVAEE DEDDDGGIMM RSKETSSPGT DDVFTPAPSD SPSSQRIQRC LSDPGPHPEP GEGEPFFPKG Q

#### A1-LCD

MASASSSQRG RSGSGNFGGG RGGGFGGNDN FGRGGNFSGR GGFGGSRGGG GYGGSGDGYN GFGNDGSNFG GGGSYNDFGN YNNQSSNFGP MKGGNFGGRS SGGSGGGGQY FAKPRNQGGY GGSSSSSSYG SGRRF

#### *α*-Synuclein

MDVFMKGLSK AKEGVVAAAE KTKQGVAEAA GKTKEGVLYV GSKTKEGVVH GVATVAEKTK EQVTNVGGAV VTGVTAVAQK TVEGAGSIAA ATGFVKKDQL GKNEEGAPQE GILEDMPVDP DNEAYEMPSE EGYQDYEPEA

#### K17

MSSPGSPGTP GSRSRTPSLP TPPTREPKKV AVVRTPPKSP SSAKSRLQTA PVPMPDLKNV KSKIGSTENL KHQPGGGKVQ IVYKPVDLSK VTSKCGSLGN IHHKPGGGQV EVKSEKLDFK DRVQSKIGSL DNITHVPGGG NKKIE

#### N98

GCFNKSFGTP FGGGTGGFGT TSTFGQNTGF GTTSGGAFGT SAFGSSNNTG GLFGNSQTKP GGLFGTSSFS QPATSTSTGF GFGTSTGTAN TLFGTASTGT SLFSSQNNAF AQNKPTGFGN FGTSTSSGGL FGTTNTTSNP FGSTSGSLFG P

#### K27

MSSPGSPGTP GSRSRTPSLP TPPTREPKKV AVVRTPPKSP SSAKSRLQTA PVPMPDLKNV KSKIGSTENL KHQPGGGKVQ IVYKPVDLSK VTSKCGSLGN IHHKPGGGQV EVKSEKLDFK DRVQSKIGSL DNITHVPGGG NKKIETHKLT FRENAKAKTD HGAEIVY

#### K10

MQTAPVPMPD LKNVKSKIGS TENLKHQPGG GKVQIVYKPV DLSKVTSKCG SLGNIHHKPG GGQVEVKSEK LDFKDRVQSK IGSLDNITHV PGGGNKKIET HKLTFRENAK AKTDHGAEIV YKSPVVSGDT SPRHLSNVSS TGSIDMVDSP QLATLADEVS ASLAKQGL

#### K16

MSSPGSPGTP GSRSRTPSLP TPPTREPKKV AVVRTPPKSP SSAKSRLQTA PVPMPDLKNV KSKIGSTENL KHQPGGGKVQ IINKKLDLSN VQSKCGSKDN IKHVPGGGSV QIVYKPVDLS KVTSKCGSLG NIHHKPGGGQ VEVKSEKLDF KDRVQSKIGS LDNITHVPGG GNKKIE

#### NSP

MAEPRQEFEV MEDHAGTYGL GDRKDQGGYT MHQDQEGDTD AGLKAEEAGI GDTPSLEDEA AGHVTQARMV SKSKDGTGSD DKKAKGADGK TKIATPRGAA PPGQKGQANA TRIPAKTPPA PKTPPSSGEP PKSGDRSGYS SPGSPGTPGS RSRTPSLPTP PTREPKKVAV VRTPPKSPSS AKSRL

#### K25

MAEPRQEFEV MEDHAGTYGL GDRKDQGGYT MHQDQEGDTD AGLKAEEAGI GDTPSLEDEA AGHVTQARMV SKSKDGTGSD DKKAKGADGK TKIATPRGAA PPGQKGQANA TRIPAKTPPA PKTPPSSGEP PKSGDRSGYS SPGSPGTPGS RSRTPSLPTP PTREPKKVAV VRTPPKSPSS AKSRL

#### K32

MSSPGSPGTP GSRSRTPSLP TPPTREPKKV AVVRTPPKSP SSAKSRLQTA PVPMPDLKNV KSKIGSTENL KHQPGGGKVQ IINKKLDLSN VQSKCGSKDN IKHVPGGGSV QIVYKPVDLS KVTSKCGSLG NIHHKPGGGQ VEVKSEKLDF KDRVQSKIGS LDNITHVPGG GNKKIETHKL TFRENAKAKT DHGAEIVY

### The sequence of Multi-Domain Proteins

#### hnRNPA1

MSKSESPKEP EQLRKLFIGG LSFETTDESL RSHFEQWGTL TDCVVMRDPN TKRSRGFGFV TYATVEEVDA AMNARPHKVD GRVVEPKRAV SREDSQRPGA HLTVKKIFVG GIKEDTEEHH LRDYFEQYGK IEVIEIMTDR GSGKKRGFAF VTFDDHDSVD KIVIQKYHTV NGHNCEVRKA LSKQEMASAS SSQRGRSGSG NFGGGRGGGF GGNDNFGRGG NFSGRGGFGG SRGGGGYGGS GDGYNGFGND GGYGGGGPGY SGGSRGYGSG GQGYGNQGSG YGGSGSYDSY NNGGGGGFGG GSGSNFGGGG SYNDFGNYNN QSSNFGPMKG GNFGGRSSGP YGGGGQYFAK PRNQGGYGGS SSSSSYGSGR RF

#### TDP-43

MSEYIRVTED ENDEPIEIPS EDDGTVLLST VTAQFPGACG LRYRNPVSQC MRGVRLVEGI LHAPDAGWGN LVYVVNYPKD NKRKMDETDA SSAVKVKRAV QKTSDLIVLG LPWKTTEQDL KEYFSTFGEV LMVQVKKDLK TGHSKGFGFV RFTEYETQVK VMSQRHMIDG RWCDCKLPNS KQSQDEPLRS RKVFVGRCTE DMTEDELREF FSQYGDVMDV FIPKPFRAFA FVTFADDQIA QSLCGEDLII KGISVHISNA EPKHNSNRQL ERSGRFGGNP GGFGNQGGFG NSRGGGAGLG NNQGSNMGGG MNFGAFSINP AMMAAAQAAL QSSWGMMGML ASQQNQSGPS GNNQNQGNMQ REPNQAFGSG NNSYSGSNSG AAIGWGSASN AGSGSGFNGG FGSSMDSKSS GWGM

#### G3BP1

MVMEKPSPLL VGREFVRQYY TLLNQAPDML HRFYGKNSSY VHGGLDSNGK PADAVYGQKE IHRKVMSQNF TNCHTKIRHV DAHATLNDGV VVQVMGLLSN NSQALRRFMQ TFVLAPEGSV ANKFYVHNDI FRYQDEVFGG FVTEPQEESE EEVEEPEERQ QTPEVVPDDS GTFYDQAVVS NDMEEHLEEP VAEPEPDPEP EPEQEPVSEI QEEKPEPVLE ETAPEDAQKS SSPAPADIAQ TVQEDLRTFS WASVTSKNLP SGAVPVTGIP PHVVKVPASQ PRPESKPESQ IPPQRPQRDQ RVREQRINIP PQRGPRPIRE AGEQGDIEPR RMVRHPDSHQ LFIGNLPHEV DKSELKDFFQ SYGNVVELRI NSGGKLPNFG FVVFDDSEPV QKVLSNRPIM FRGEVRLNVE EKKTRAAREG DRRDNRLRGP GGPRGGLGGG MRGPPRGGMV QKPGFGVGRG LAPRQ

#### FUS

MASNDYTQQA TQSYGAYPTQ PGQGYSQQSS QPYGQQSYSG YSQSTDTSGY GQSSYSSYGQ SQNTGYGTQS TPQGYGSTGG YGSSQSSQSS YGQQSSYPGY GQQPAPSSTS GSYGSSSQSS SYGQPQSGSY SQQPSYGGQQ QSYGQQQSYN PPQGYGQQNQ YNSSSGGGGG GGGGGNYGQD QSSMSSGGGS GGGYGNQDQS GGGGSGGYGQ QDRGGRGRGG SGGGGGGGGG GYNRSSGGYE PRGRGGGRGG RGGMGGSDRG GFNKFGGPRD QGSRHDSEQD NSDNNTIFVQ GLGENVTIES VADYFKQIGI IKTNKKTGQP MINLYTDRET GKLKGEATVS FDDPPSAKAA IDWFDGKEFS GNPIKVSFAT RRADFNRGGG NGRGGRGRGG PMGRGGYGGG GSGGGGRGGF PSGGGGGGGQ QRAGDWKCPN PTCENMNFSW RNECNQCKAP KPDGPGGGPG GSHMGGNYGD DRRGGRGGYD RGGYRGRGGD RGGFRGGRGG GDRGGFGPGK MDSRGEHRQD RRERPY

##### Comparison of epsilon of SOP-IDP and SOP-Multi-*π* models for *π* −*π* and cation−*π* interaction pairs

**Figure S1:**
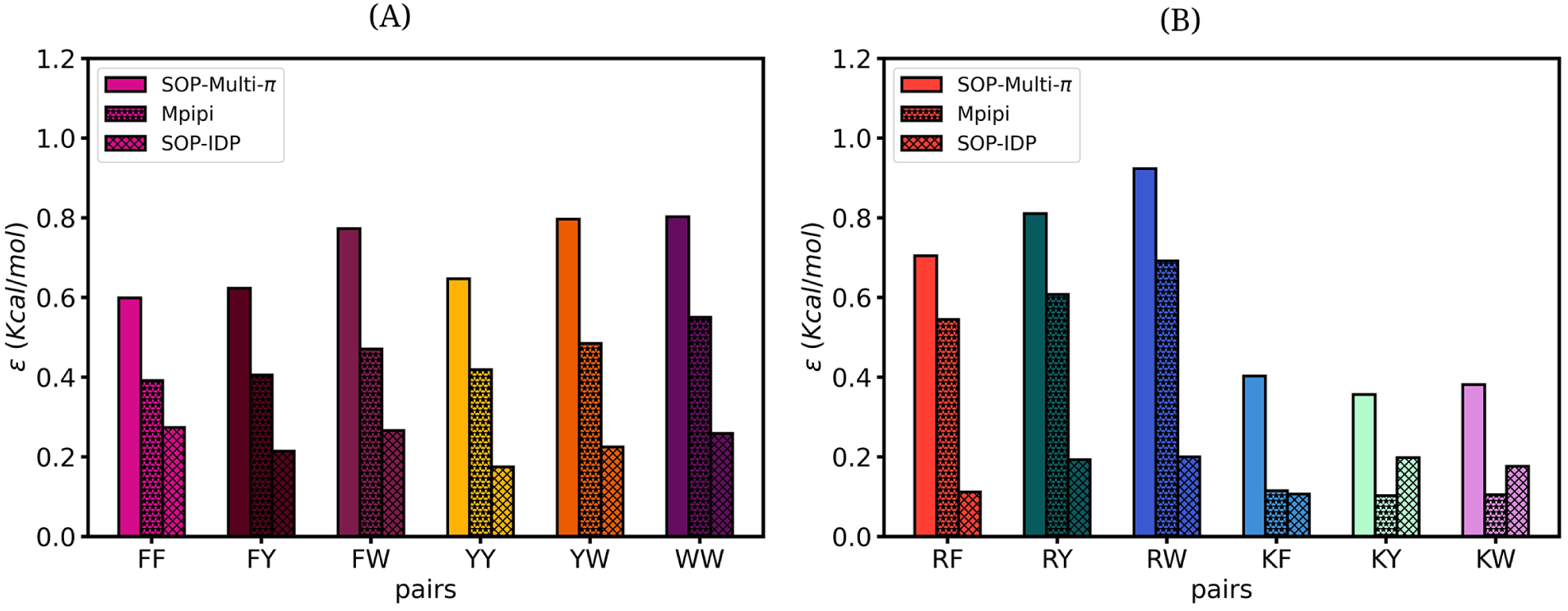
The bar plot to show that ordering of *π*−*π* interaction (A) and cation-*π* interaction (B) of the SOP-Multi-*π*, Mpipi, and SOP-IDP models.

**Figure S2:**
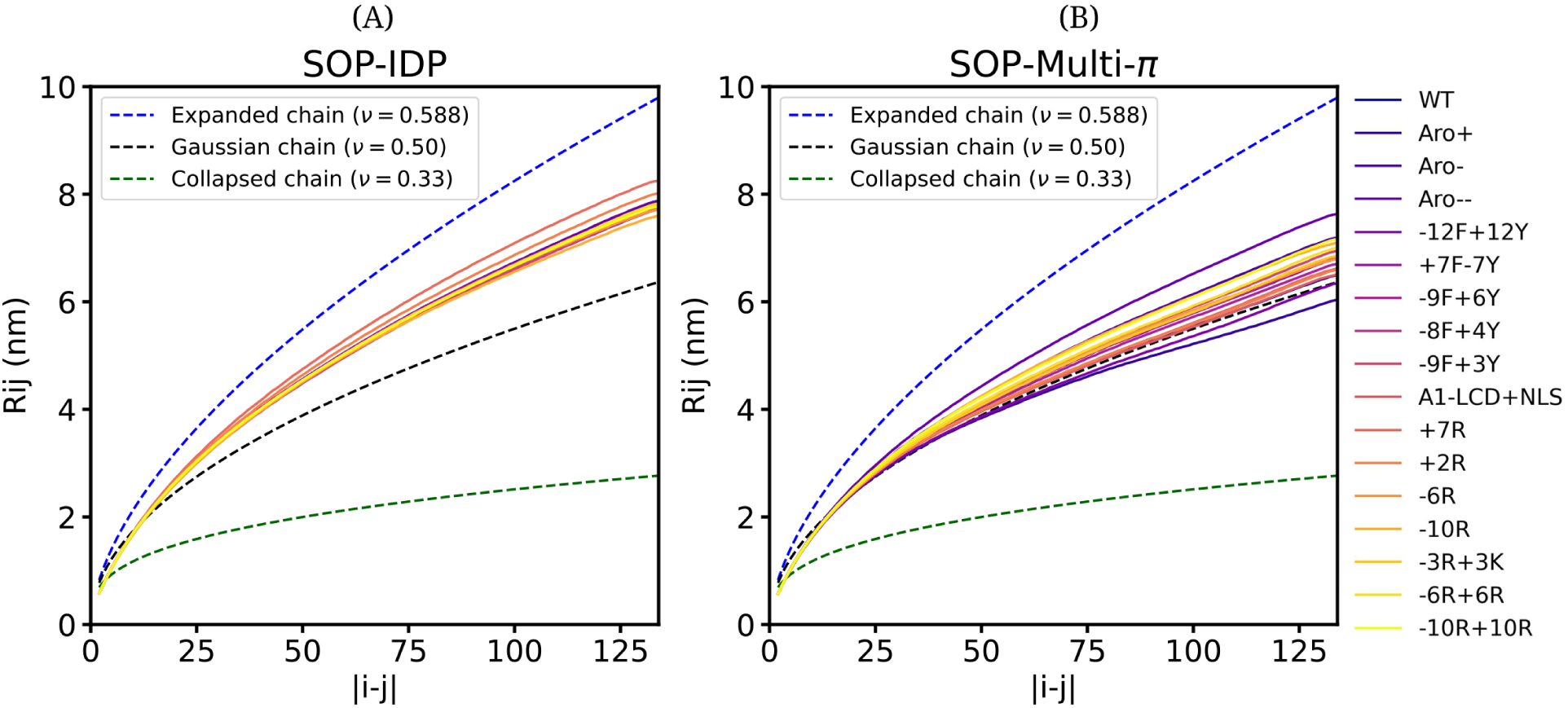
Flory exponent of A1-LCD and its Variants for the SOP-IDP and SOP-Multi-*π* models.

**Figure S3:**
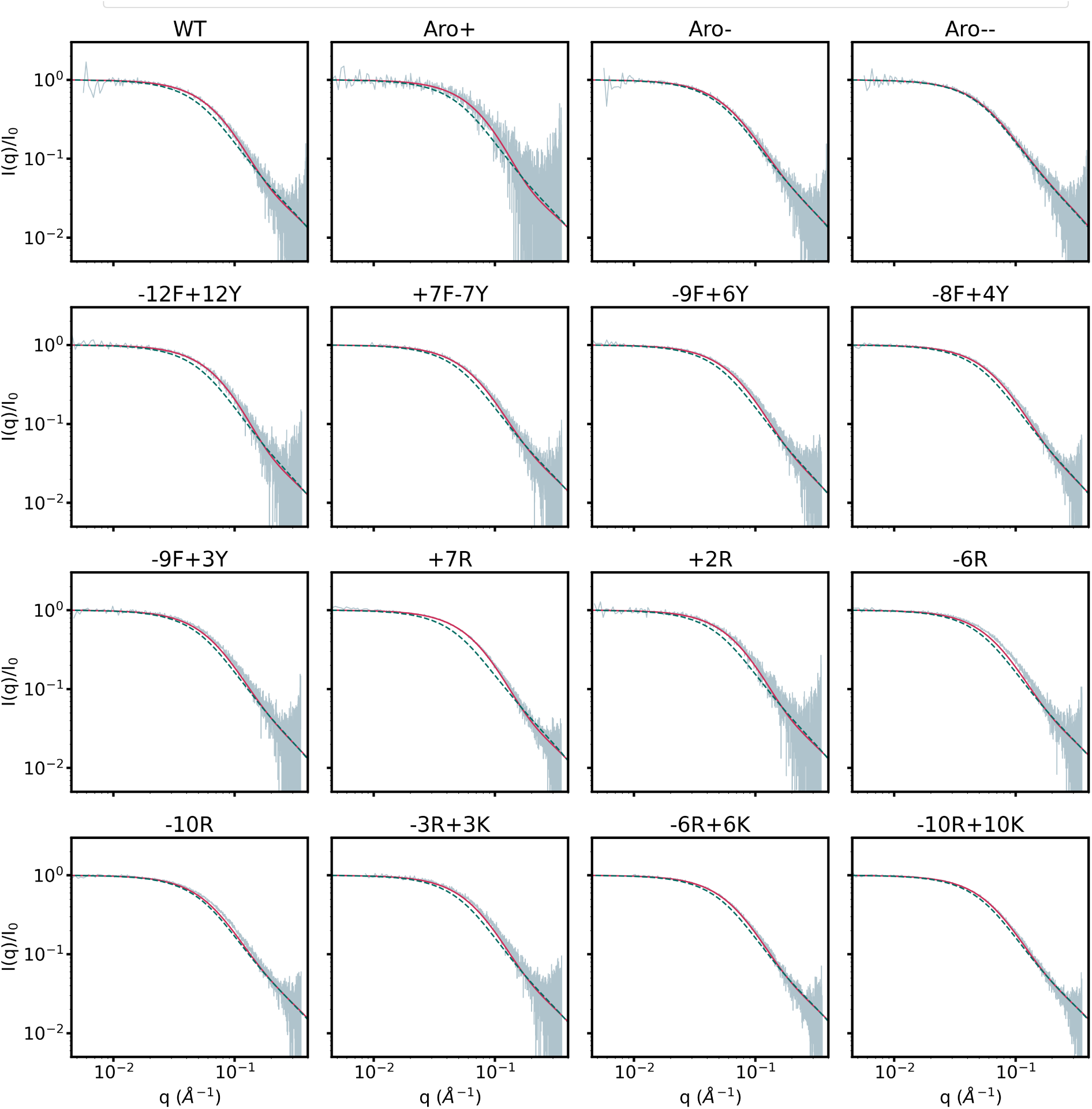
SAXS curve, grey colour curve is the experimental data, green colour dash curve is the SOP-IDP simulation result, and pink colour curve is the SOP-Multi-*π* result.

**Table S1:** Table contains the one-letter and three-letter codes of the amino acids in the 1*^st^* and 2*^nd^* columns. The van der Waals radius, charge, type, and mass of the amino acid residues are tabulated in the 4*^th^*, 5*^th^*, 6*^th^*, and 7*^th^* columns, respectively.^3^

| 3-Letter | 1-Letter | ID | vdW radius (Å) | Charge(e) | Type | Mass (amu) |
| --- | --- | --- | --- | --- | --- | --- |
| $C_\alpha$-atom | BB | 1 | 1.90 | 0 | $C_\alpha$ | 56.044 |
| GLY | G | 2 | 2.25 | 0 | SpCase | 57.051 |
| ALA | A | 3 | 2.52 | 0 | HPhobic | 15.034 |
| ARG | R | 4 | 3.28 | 1 | Charged | 101.150 |
| LYS | K | 5 | 3.18 | 1 | Charged | 73.136 |
| HIS | H | 6 | 3.04 | 0/1 | Charged | 81.095 |
| ASP | D | 7 | 2.79 | -1 | Charged | 58.035 |
| GLU | E | 8 | 2.96 | -1 | Charged | 72.062 |
| SER | S | 9 | 2.59 | 0 | Polar | 31.033 |
| THR | T | 10 | 2.81 | 0 | Polar | 45.060 |
| ASN | N | 11 | 2.84 | 0 | Polar | 58.035 |
| GLN | Q | 12 | 3.01 | 0 | Polar | 61.080 |
| CYS | C | 13 | 2.74 | 0 | SpCase | 47.099 |
| PRO | P | 14 | 2.78 | 0 | SpCase | 41.071 |
| ILE | I | 15 | 3.09 | 0 | Hphobic | 57.114 |
| LEU | L | 16 | 3.09 | 0 | Hphobic | 57.114 |
| MET | M | 17 | 3.09 | 0 | Hphobic | 75.152 |
| PHE | F | 18 | 3.18 | 0 | Aromatic | 91.130 |
| TRP | W | 19 | 3.39 | 0 | Aromatic | 130.166 |
| TYR | Y | 20 | 3.23 | 0 | Aromatic | 107.129 |
| VAL | V | 21 | 2.93 | 0 | HPhobic | 43.087 |

**Table S2:** Table of sequence properties of single-chain A1-LCD and its variant used in rescaling SOP-Multi-*π* force field. The f_+_, f*_−_*, FCR, NCPR, and f*_Aro_* denote fraction of positive charge, fraction of negative charge, fraction of charged residue, Net charge per residue, and fraction of aromatic residue, respectively. Trp, Phe, Tyr, Arg, and Lys denote the number of aromatic (Trp, Phe, and Tyr) and positive charge (Arg and Lys) residues present in the sequence of IDPs.

| IDPs | $f_+$ | $f_-$ | FCR | NCPR | $f_{Aro}$ | Trp | Phe | Tyr | Arg | Lys |
| --- | --- | --- | --- | --- | --- | --- | --- | --- | --- | --- |
| A1-LCD | 0.09 | 0.03 | 0.12 | 0.06 | 0.14 | 0 | 12 | 7 | 10 | 2 |
| Aro+ | 0.09 | 0.03 | 0.12 | 0.06 | 0.18 | 0 | 15 | 9 | 10 | 2 |
| Aro- | 0.09 | 0.03 | 0.12 | 0.06 | 0.10 | 0 | 8 | 5 | 10 | 2 |
| Aro-- | 0.09 | 0.03 | 0.12 | 0.06 | 0.04 | 0 | 4 | 2 | 10 | 2 |
| -12F+12Y | 0.09 | 0.03 | 0.12 | 0.06 | 0.14 | 0 | 0 | 19 | 10 | 2 |
| +7F-7Y | 0.09 | 0.03 | 0.12 | 0.06 | 0.14 | 0 | 19 | 0 | 10 | 2 |
| -9F+6Y | 0.09 | 0.03 | 0.12 | 0.06 | 0.12 | 0 | 3 | 13 | 10 | 2 |
| -8F+4Y | 0.09 | 0.03 | 0.12 | 0.06 | 0.11 | 0 | 4 | 11 | 10 | 2 |
| -9F+3Y | 0.09 | 0.03 | 0.12 | 0.06 | 0.10 | 0 | 3 | 10 | 10 | 2 |
| A1-LCD<sup>+NLS</sup> | 0.09 | 0.03 | 0.12 | 0.06 | 0.15 | 0 | 12 | 8 | 10 | 2 |
| +7R | 0.14 | 0.03 | 0.17 | 0.11 | 0.15 | 0 | 12 | 8 | 17 | 2 |
| +2R | 0.10 | 0.03 | 0.13 | 0.07 | 0.15 | 0 | 12 | 8 | 12 | 2 |
| -6R | 0.04 | 0.03 | 0.07 | 0.01 | 0.15 | 0 | 12 | 8 | 4 | 2 |
| -10R | 0.01 | 0.03 | 0.04 | 0.01 | 0.15 | 0 | 12 | 8 | 0 | 2 |
| -3R+3K | 0.09 | 0.03 | 0.12 | 0.06 | 0.14 | 0 | 12 | 7 | 7 | 5 |
| -6R+6K | 0.09 | 0.03 | 0.12 | 0.06 | 0.14 | 0 | 12 | 7 | 4 | 8 |
| -10R+10K | 0.09 | 0.03 | 0.12 | 0.06 | 0.14 | 0 | 12 | 7 | 0 | 12 |

**Table S3:**
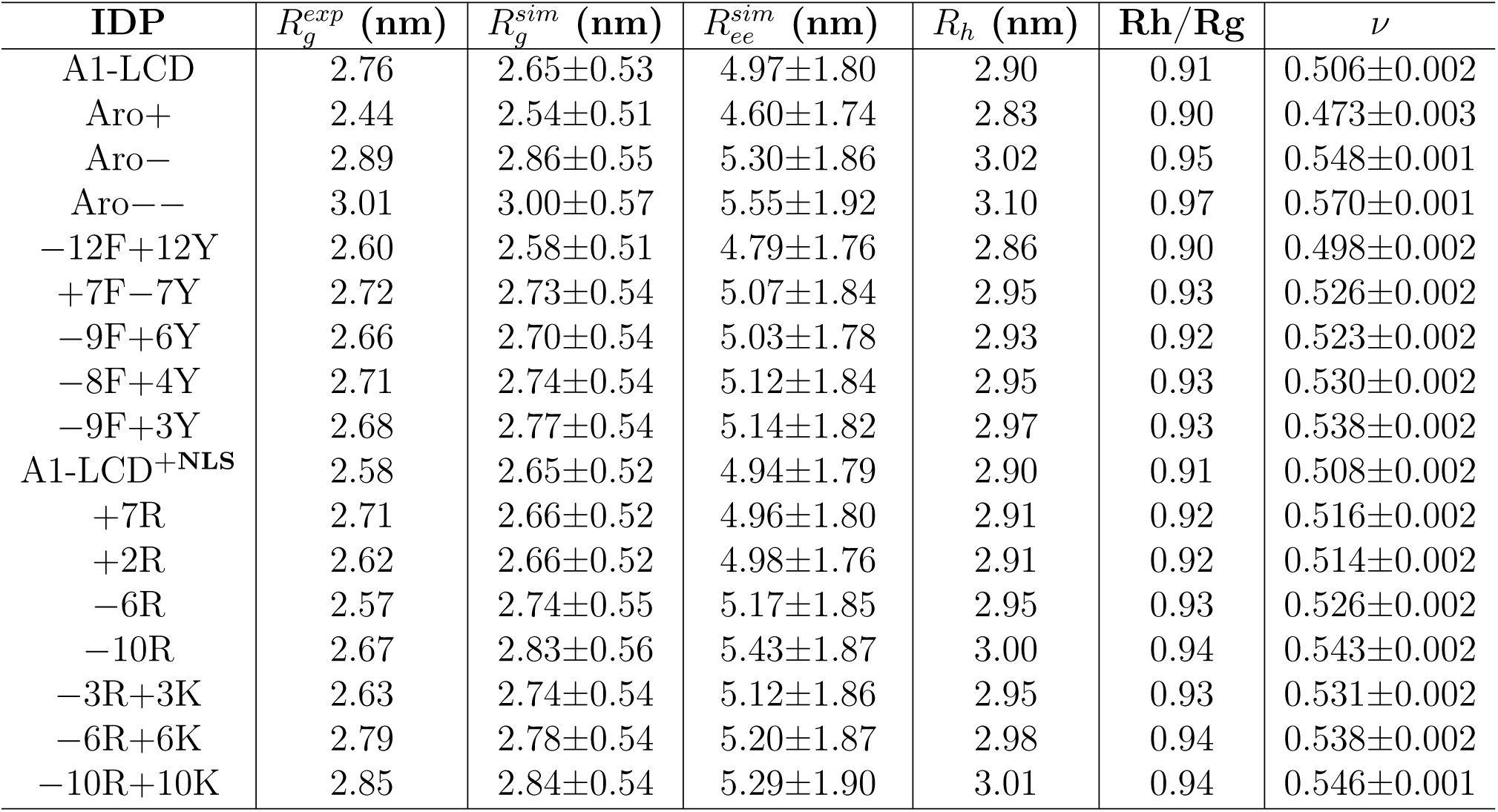
Table of A1-LCD (WT) and its variants studies in our work to rescale the SOP-Multi-*π* force field. Where 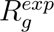 and 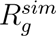 are experimental and simulated radius of gyration. 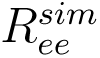, *R_h_*, and *ν* are the simulated end-to-end distance, hydrodynamic radius, and Flory exponent, respectively.

| IDP | $R_g^{exp}$ (nm) | $R_g^{sim}$ (nm) | $R_{ee}^{sim}$ (nm) | $R_h$ (nm) | Rh/Rg | $\nu$ |
| --- | --- | --- | --- | --- | --- | --- |
| A1-LCD | 2.76 | 2.65±0.53 | 4.97±1.80 | 2.90 | 0.91 | 0.506±0.002 |
| Aro+ | 2.44 | 2.54±0.51 | 4.60±1.74 | 2.83 | 0.90 | 0.473±0.003 |
| Aro- | 2.89 | 2.86±0.55 | 5.30±1.86 | 3.02 | 0.95 | 0.548±0.001 |
| Aro-- | 3.01 | 3.00±0.57 | 5.55±1.92 | 3.10 | 0.97 | 0.570±0.001 |
| -12F+12Y | 2.60 | 2.58±0.51 | 4.79±1.76 | 2.86 | 0.90 | 0.498±0.002 |
| +7F-7Y | 2.72 | 2.73±0.54 | 5.07±1.84 | 2.95 | 0.93 | 0.526±0.002 |
| -9F+6Y | 2.66 | 2.70±0.54 | 5.03±1.78 | 2.93 | 0.92 | 0.523±0.002 |
| -8F+4Y | 2.71 | 2.74±0.54 | 5.12±1.84 | 2.95 | 0.93 | 0.530±0.002 |
| -9F+3Y | 2.68 | 2.77±0.54 | 5.14±1.82 | 2.97 | 0.93 | 0.538±0.002 |
| A1-LCD<sup>+NLS</sup> | 2.58 | 2.65±0.52 | 4.94±1.79 | 2.90 | 0.91 | 0.508±0.002 |
| +7R | 2.71 | 2.66±0.52 | 4.96±1.80 | 2.91 | 0.92 | 0.516±0.002 |
| +2R | 2.62 | 2.66±0.52 | 4.98±1.76 | 2.91 | 0.92 | 0.514±0.002 |
| -6R | 2.57 | 2.74±0.55 | 5.17±1.85 | 2.95 | 0.93 | 0.526±0.002 |
| -10R | 2.67 | 2.83±0.56 | 5.43±1.87 | 3.00 | 0.94 | 0.543±0.002 |
| -3R+3K | 2.63 | 2.74±0.54 | 5.12±1.86 | 2.95 | 0.93 | 0.531±0.002 |
| -6R+6K | 2.79 | 2.78±0.54 | 5.20±1.87 | 2.98 | 0.94 | 0.538±0.002 |
| -10R+10K | 2.85 | 2.84±0.54 | 5.29±1.90 | 3.01 | 0.94 | 0.546±0.001 |

**Table S4:** List of IDPs used to validate the SOP-Multi-*π* forcefield. K10, K17, K19, K25, and K27 are the truncated hTau23 protein and K16, K18, and K32 are the truncated hTau40 protein. Where 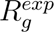 and 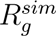 are experimental and simulated radius of gyration. 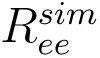, *R_h_* and *ν* are the simulated end-to-end distance, hydrodynamic radius, and Flory exponent, respectively.

| IDP | $R_g^{exp}$ (nm) | $R_g^{sim}$ (nm) | $R_{ee}^{sim}$ (nm) | $R_h$ (nm) | Rh/Rg | $\nu$ |
| --- | --- | --- | --- | --- | --- | --- |
| Histatin5 | 1.35 | 1.22±0.15 | 1.83±0.46 | 1.43 | 0.86 | 0.559±0.007 |
| RS-repeat | 1.26 | 1.20±0.15 | 1.94±0.50 | 1.42 | 0.85 | 0.580±0.004 |
| N49 | 1.69 | 1.50±0.21 | 2.52±0.65 | 1.68 | 0.89 | 0.590±0.003 |
| NLS | 2.33 | 1.98±0.25 | 3.08±0.76 | 1.95 | 1.02 | 0.653±0.003 |
| Histatin5-2 | 1.87 | 1.90±0.28 | 3.01±0.89 | 1.97 | 0.97 | 0.600±0.004 |
| ACTR | 2.63 | 2.28±0.37 | 3.85±1.15 | 2.34 | 0.97 | 0.576±0.002 |
| NUS | 2.41 | 2.42±0.42 | 4.15±1.31 | 2.48 | 0.98 | 0.576±0.002 |
| sfAFP | 2.31 | 2.18±0.41 | 4.14±1.38 | 2.37 | 0.92 | 0.551±0.002 |
| Nucleoporin | 2.70 | 2.43±0.42 | 4.15±1.29 | 2.49 | 0.98 | 0.573±0.002 |
| Ash1 | 2.85 | 2.65±0.44 | 4.34±1.33 | 2.60 | 1.02 | 0.601±0.002 |
| SH4-UD | 2.86 | 2.53±0.44 | 4.43±1.39 | 2.56 | 0.99 | 0.581±0.002 |
| Sic1 | 3.21 | 2.75±0.47 | 4.60±1.40 | 2.70 | 1.02 | 0.591±0.002 |
| P53 | 2.87 | 2.72±0.48 | 4.54±1.43 | 2.71 | 1.00 | 0.572±0.002 |
| IBB | 3.12 | 2.88±0.52 | 5.18±1.53 | 2.81 | 1.03 | 0.582±0.001 |
| ColNT | 2.83 | 2.58±0.47 | 4.53±1.51 | 2.68 | 0.97 | 0.558±0.001 |
| K19 | 3.50 | 2.89±0.49 | 4.87±1.56 | 2.83 | 1.02 | 0.588±0.002 |
| Prothymosin-α | 3.78 | 3.20±0.59 | 5.38±1.77 | 3.05 | 1.05 | 0.546±0.005 |
| p15PAF | 2.81 | 3.29±0.55 | 5.42±1.65 | 3.09 | 1.06 | 0.609±0.002 |
| NUL1 | 3.50 | 2.93±0.54 | 4.99±1.70 | 2.94 | 1.00 | 0.559±0.002 |
| hNL3cyt | 3.00 | 3.11±0.58 | 5.28±1.80 | 3.07 | 1.02 | 0.560±0.001 |
| ERM-TADn | 3.96 | 3.12±0.58 | 4.92±1.70 | 3.09 | 1.01 | 0.552±0.002 |
| RNaseA | 3.30 | 3.29±0.60 | 5.66±1.75 | 3.18 | 1.04 | 0.572±0.002 |
| K18 | 3.80 | 3.40±0.61 | 5.80±1.91 | 3.27 | 1.04 | 0.584±0.002 |
| hNHE1cdt | 3.75 | 3.31±0.61 | 5.65±1.85 | 3.23 | 1.02 | 0.569±0.001 |
| A1-LCD | 2.76 | 2.65±0.53 | 4.96±1.80 | 2.90 | 0.91 | 0.497±0.001 |
| α-Synuclein | 3.50 | 3.63±0.66 | 6.26±2.02 | 3.43 | 1.06 | 0.593±0.002 |
| K17 | 3.60 | 3.76±0.66 | 6.43±1.98 | 3.53 | 1.07 | 0.599±0.002 |
| N98 | 2.86 | 3.30±0.65 | 5.79±2.08 | 3.34 | 0.99 | 0.541±0.002 |
| K27 | 3.70 | 4.13±0.75 | 6.98±2.26 | 3.83 | 1.08 | 0.597±0.002 |
| K10 | 4.00 | 3.93±0.74 | 6.89±2.24 | 3.74 | 1.05 | 0.568±0.002 |
| K16 | 3.90 | 4.19±0.76 | 7.26±2.32 | 3.91 | 1.07 | 0.591±0.001 |
| NSP | 4.10 | 3.81±0.75 | 6.49±2.34 | 3.72 | 1.02 | 0.551±0.001 |
| K25 | 4.10 | 4.12±0.80 | 6.85±2.32 | 3.92 | 1.05 | 0.575±0.001 |
| K32 | 4.20 | 4.52±0.82 | 7.76±2.56 | 4.18 | 1.08 | 0.587±0.001 |

## Supporting Information (User Manual)

### Building and simulating multi-component biomolecular condensates with the SOP-Multi-*π* force field

September 30, 2026

This document provides a protocol for constructing and running coarse-grained condensate simulations with the SOP-Multi-*π* force field in LAMMPS. It covers systems containing a single protein species and systems containing two, three or more distinct species, including multidomain proteins with both folded domains and intrinsically disordered regions (IDRs). The protocol describes the construction workflow and provides references for the scripts and the underlying force-field framework. The underlying SOP-MULTI coarse-grained model on which this force field is built is described by Baratam and Srivastava [1].

The protocol assumes LAMMPS built with the SOP-MULTI package, which provides atom_style *→* sop_bead, pair_style lj/sop, pair_style debye/sop and `special_bonds sop_ff`.

#### 1 Overview of the workflow

System construction proceeds in five stages. The first two stages are common to all systems, whereas stages 3–5 depend on whether the system contains a single species or multiple species.

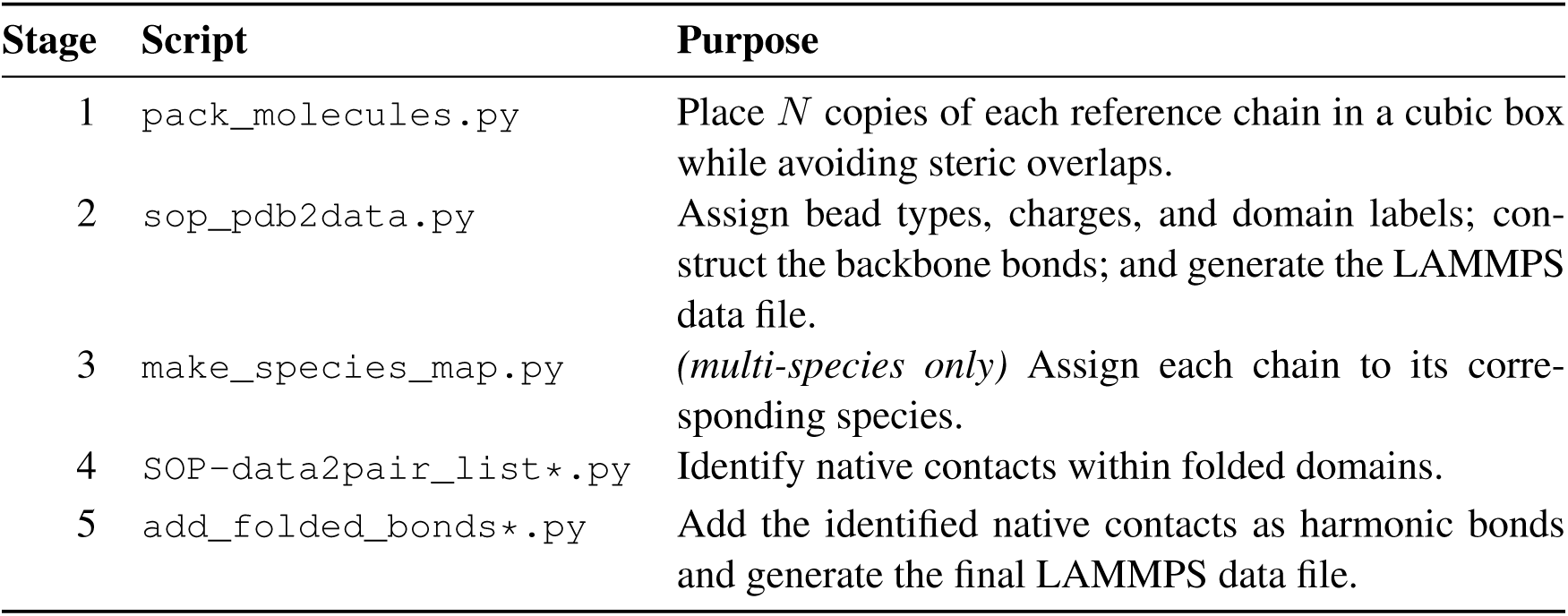

| Stage | Script | Purpose |
| --- | --- | --- |
| 1 | `pack_molecules.py` | Place $N$ copies of each reference chain in a cubic box while avoiding steric overlaps. |
| 2 | `sop_pdb2data.py` | Assign bead types, charges, and domain labels; construct the backbone bonds; and generate the LAMMPS data file. |
| 3 | `make_species_map.py` | *(multi-species only)* Assign each chain to its corresponding species. |
| 4 | `SOP-data2pair_list*.py` | Identify native contacts within folded domains. |
| 5 | `add_folded_bonds*.py` | Add the identified native contacts as harmonic bonds and generate the final LAMMPS data file. |

The force-field file and the LAMMPS input scripts are independent of the number of chains and species in the system. All proteins use the same 21-entry amino-acid bead library, resulting in 42 atom types in the current implementation (21 disordered + 21 folded copies). The `read_data` command automatically obtains the atom count, bond count, and simulation-box dimensions from the data file. Consequently, the LAMMPS input scripts can be reused for systems with different numbers of chains and species; only the `data_filename` variable needs to be changed.

##### 1.1 Required files

| File | Description |
| --- | --- |
| `<NAME>-CG.pdb` | Coarse-grained reference structure, one chain, two beads per residue (backbone $C\alpha$; side-chain bead at the side-chain centre of mass; glycine backbone only). One per species. |
| `domain_data-<NAME>.dat` | Folded-domain residue ranges, one row per range as `from to domain_id`. One file per species; use `none` for a fully disordered chain. |
| `sop_model_library/sop_bead_types.dat` | Bead library. **Must contain exactly 21 entries**; `sop_pdb2data.py` generates the 21 folded copies itself, so a manually expanded library is doubled twice. Shared by all species. |
| `ff_sop_42.input` | Pair coefficients for all 42 types: 903 `lj/sop` pairs ($42 \times 43/2$) and 36 `debye/sop` pairs. |

##### 1.2 Building the LAMMPS executable

The construction scripts described below produce data files for a LAMMPS build that includes the SOP-MULTI package. That package supplies the `sop_bead` atom style, the `sop_ff` special-bonds setting, and the `lj/sop` and `debye/sop` pair styles used throughout this protocol [1].

The SOP-MULTI package is maintained for a specific LAMMPS release. Use the LAMMPS version specified in the SOP-MULTI repository README. Obtain the corresponding release from https://www.lammps.org/, extract it, and download the SOP-MULTI sources from https://github.com/codesrivastavalab/SOP-MULTI. Copy the contents of `lammps-SOP-MULTI-src` into the LAMMPS `src` directory.

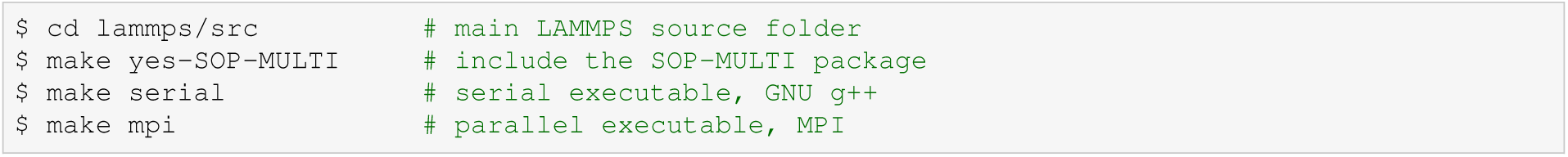

Depending on the build target the executable is named lmp_serial, lmp_mpi.

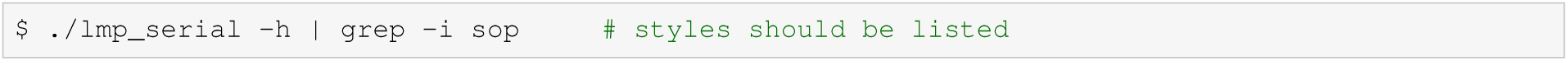

Full documentation of the individual SOP-MULTI LAMMPS features, including syntax and usage for each atom, bond and pair style, is given in the SOP-MULTI package documentation [1].

##### 1.3 Software requirements

Python 3.8 or later is required, together with NumPy and SciPy. SciPy is used only for the `cKDTree` *→* neighbour search in Stage 4. The remaining construction scripts use only the Python standard library and do not require any additional compiled dependencies or external simulation packages. Running the resulting data files requires LAMMPS built with the SOP package.

#### 2 Single-species systems

For a homogeneous system — all chains the same protein — every chain is a rigid copy of the reference structure, so folded contacts need be identified only once and can be replicated across all copies. `SOP-`*→* `data2pair_list.py` and `add_folded_bonds.py` exploit this and are the correct tools here.

##### 2.1 Commands

Example: 10 chains of FUS in a 45 nm cubic box.

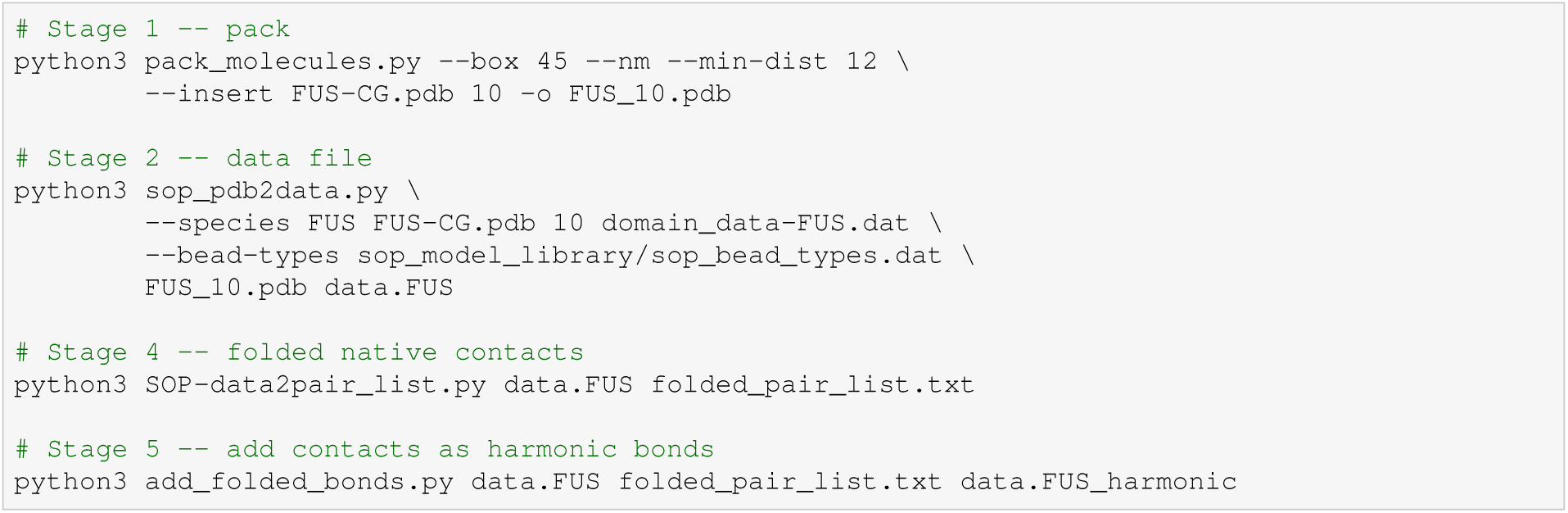

#### 3 Multi-species systems

##### 3.1 Which tools to use

`SOP-data2pair_list.py` identifies folded contacts on the first chain only, and `add_folded_bonds.` *→* `py` replicates that contact map onto every chain in the system. This is exact for a homogeneous system, but wrong for a heterogeneous one, where the fold of whichever protein comes first would be imposed on all the others. **Heterogeneous systems must therefore use the _multi tools at stages 4 and 5.** The single-species scripts detect the mismatch and exit rather than writing an incorrect data file.

Run on a homogeneous system, the multi-species tools reproduce the output of the single-species tools bond-for-bond and coefficient-for-coefficient, so they may be used for every system.

##### 3.2 Additional scripts

Contact criteria and bond coefficients are unchanged from the single-species implementation.

3.2 Additional scripts
| Script | Function |
| --- | --- |
| make_species_map.py | Writes <data>.species, mapping each molecule id to its species. The assignment is reconstructed from the insertion order, since sop_pdb2data.py does not record it. |
| SOP-data2pair_list_multi.py | Identifies folded contacts using the first chain of *each* species separately, and tags every pair with its species index. |
| add_folded_bonds_multi.py | Applies each species' contact set only to that species' chains, using the correct per-species bead offset. |

##### 3.3 Commands: two species

Example: 5 FUS + 5 G3BP1.

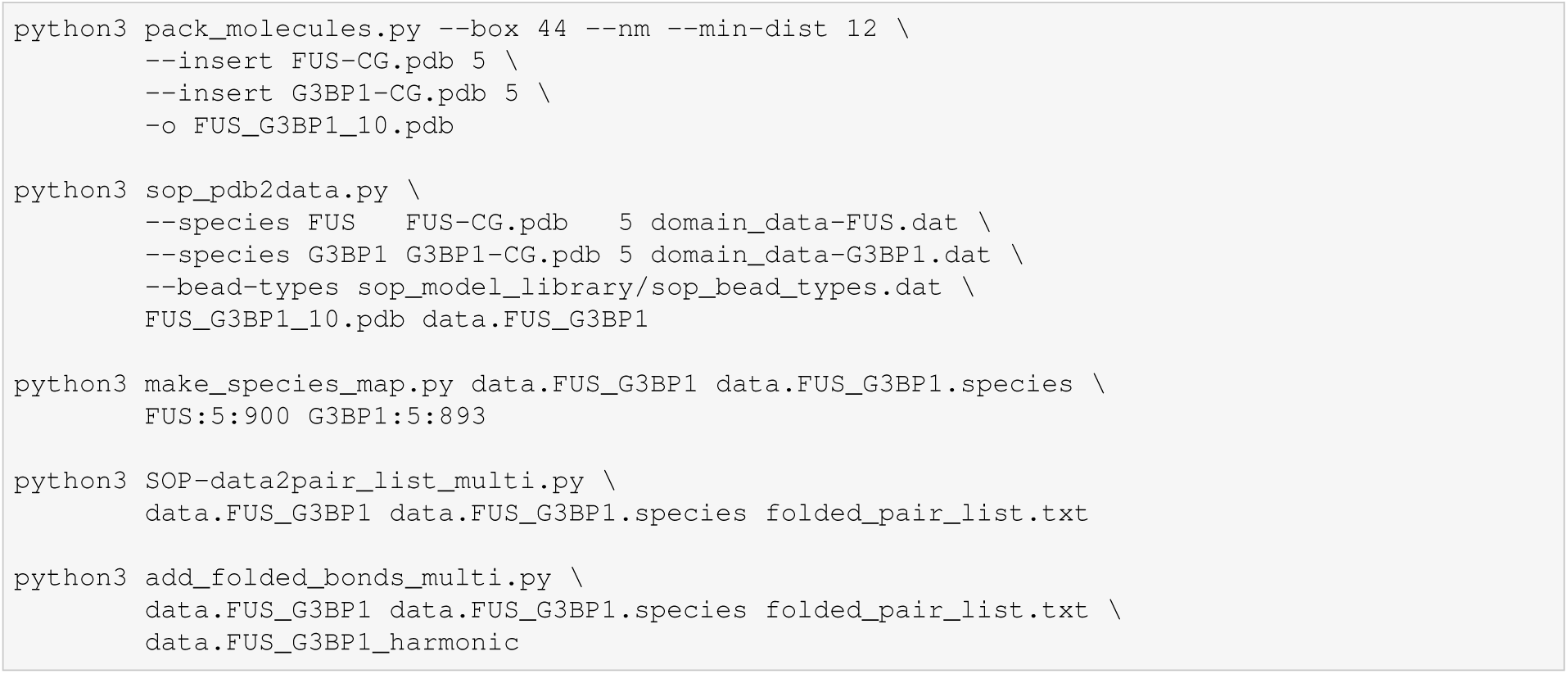

##### 3.4 Commands: three species

Example: 3 FUS + 3 G3BP1 + 4 hnRNPA1.

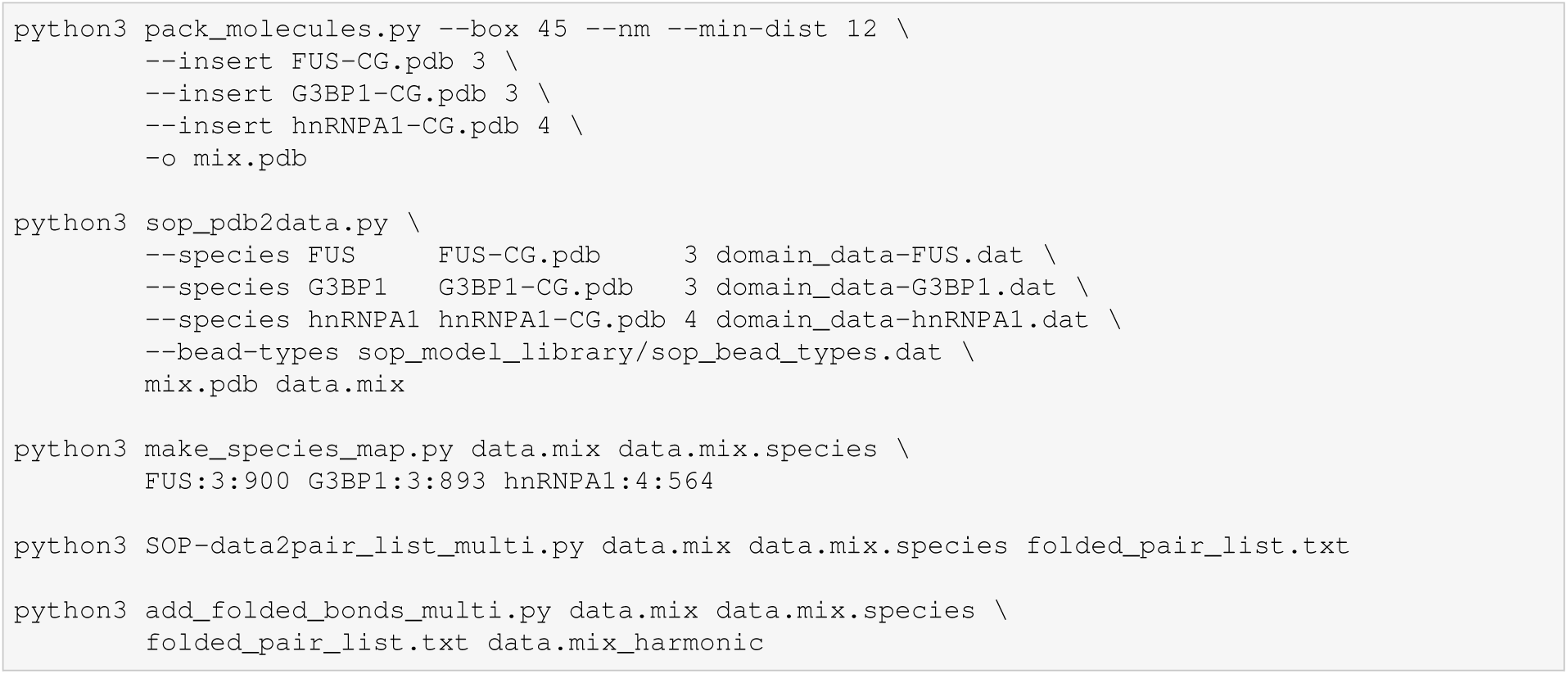

The same procedure applies to any number of species: repeat `--insert` and `--species` for each species, and provide one `NAME:NMOL:NBEADS` argument per species. **All three lists must be specified in the same order.** Molecules are packed in insertion order and residue identities are assigned by walking that same sequence, so a mismatch mislabels every bead while leaving the total atom count unchanged.

#### 4 Script reference

The commands below summarise the options required for the standard workflow.

##### 4.1 pack_molecules.py

`pack_molecules.py` packs one or more copies of reference chains into a simulation box using rigid-body placement and rejection sampling. The output is a packed PDB file containing the specified number of molecules, with a `CRYST1` record carrying the box and continuous hybrid-36 atom numbering.

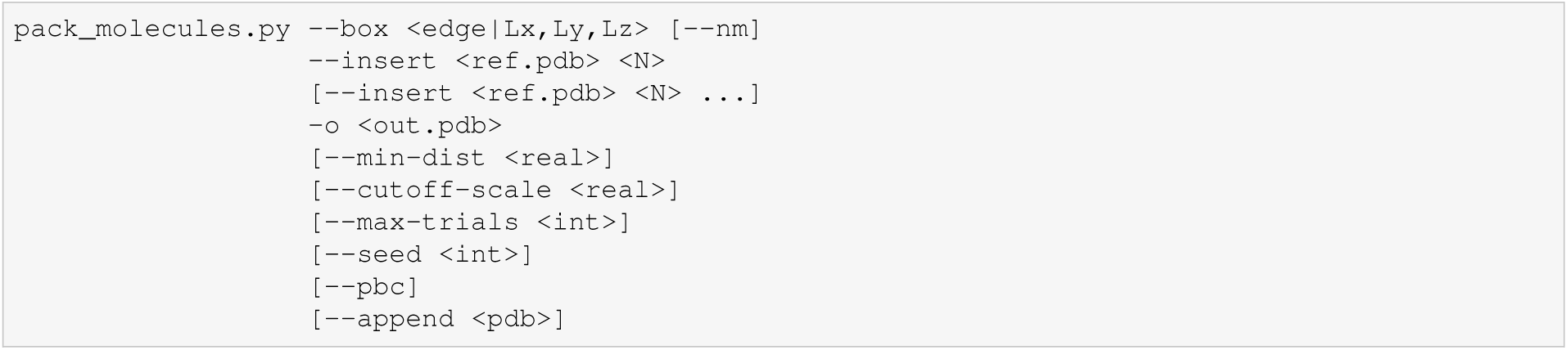

For the standard workflow we recommend `--min-dist 12`, i.e. a minimum centre-to-centre separation of 12 Å, applied as a single conservative threshold to every bead pair. All distance options (`--min-dist`, `--d-cut`, `--r0`) are in Å unless stated otherwise; force constants are in kcal mol*^−^*^1^ Å*^−^*^2^.

##### 4.2 sop_pdb2data.py

`sop_pdb2data.py` converts the packed PDB structure into a LAMMPS data file. It assigns bead types, charges, domain information and backbone bonds according to the supplied bead-type and domain-definition files. Multiple molecular species can be specified using repeated `--species` arguments, each taking a name, reference PDB, chain count and domain file, or through an INI-style configuration file.

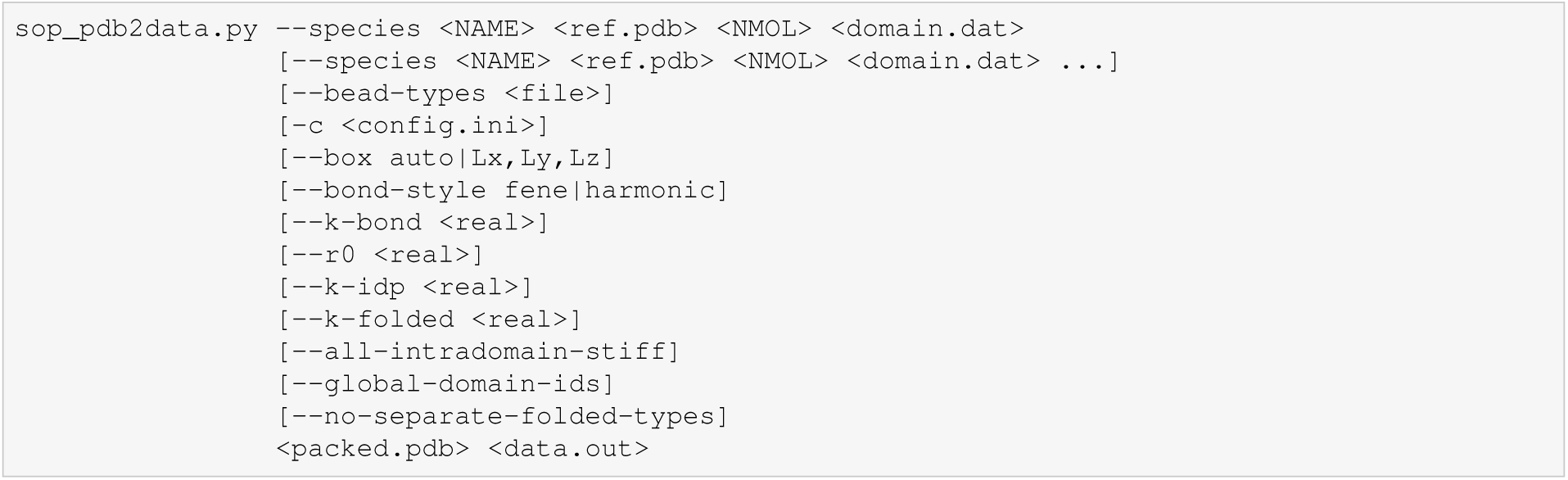

For the standard workflow, `--box auto` uses the box information stored in the packed PDB. The domain file defines folded and disordered regions; residues not explicitly assigned to a domain are treated as disordered.

##### 4.3 SOP-data2pair_list.py

`SOP-data2pair_list.py` identifies the native contacts within folded domains. A pair is written when both beads lie in the same non-zero folded domain, their separation is below *d*_cut_ = 8 Å, and their residue indices differ by more than *r*_cut_ = 2; the last condition removes the pairs already held by backbone bonds. Output columns are `atom_i`, `atom_j`, `harmonic`, *ɛ* and distance, with atom indices local to a single chain.

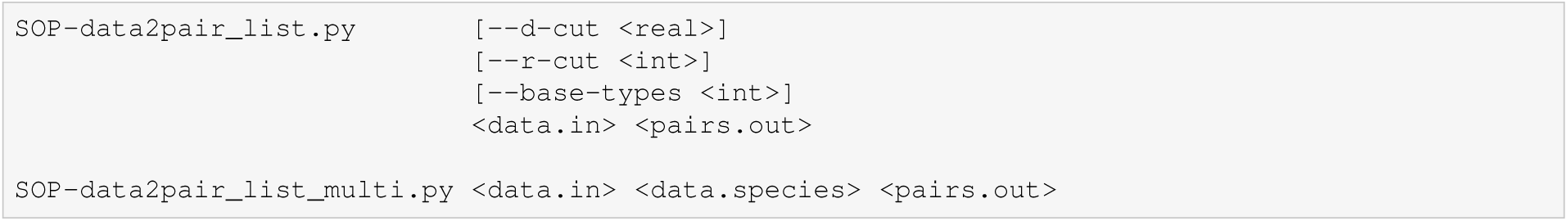

For the standard workflow, the default cutoff values are used. For multi-species systems, use `SOP-` *→* `data2pair_list_multi.py` with the corresponding species definition file.

##### 4.4 add_folded_bonds.py

`add_folded_bonds.py` combines the native-contact pair list with the LAMMPS data file and generates the final data file for simulation. The script adds the native-contact restraints and converts all bonds, backbone and native-contact alike, to harmonic form.

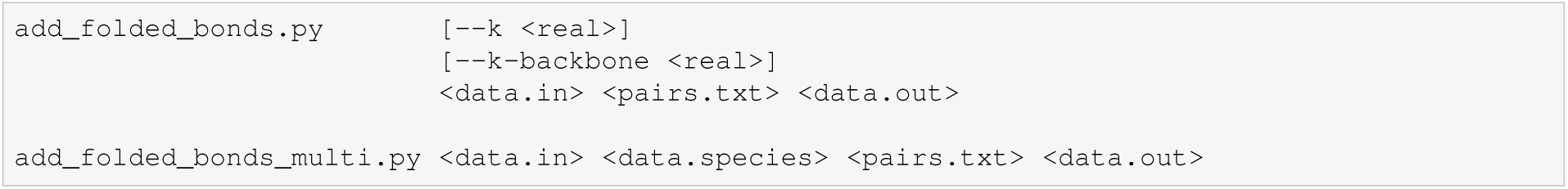

For single-species systems, use `add_folded_bonds.py`. For multi-species systems, use `add_folded_bonds_multi.py` with the corresponding species definition file.

#### 5 Force field

The simulations use the 42-type SOP force field provided in `ff_sop_42.input`, which extends the validated 21-type `forcefield.sop_multi`. Types 1–21 correspond to disordered residues and carry the rescaled *π*–*π* and cation–*π* interactions of the main text, while types 22–42 are the corresponding folded-domain types and carry the baseline values. Cross terms use the geometric mean of the two *ɛ* values.

The force field uses the following LAMMPS pair styles:

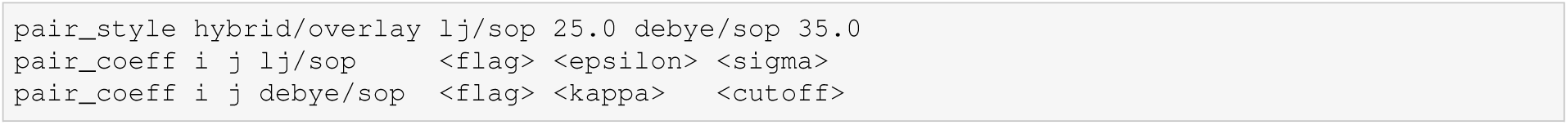

The following LAMMPS settings are required for the data files generated by the construction workflow:

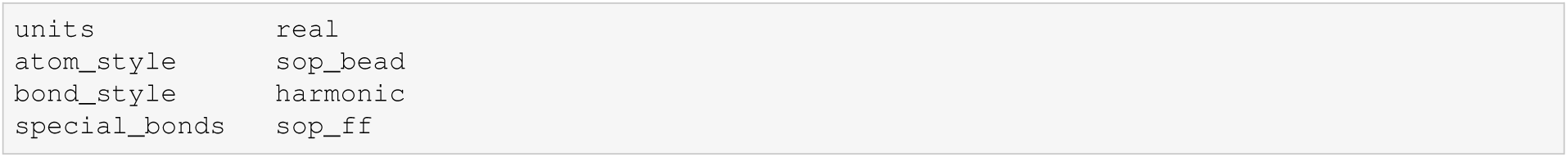

The data file does not contain a `Masses` section; masses for all 42 atom types must therefore be defined in the LAMMPS input script, with folded type *n* + 21 taking the same mass as the corresponding disordered type *n*. Charges must also be assigned in the LAMMPS input script for the charged bead types used in the model.

#### 6 Script inventory

| Script | Stage | Applies to |
| --- | --- | --- |
| `pack_molecules.py` | 1 | all systems |
| `sop_pdb2data.py` | 2 | all systems |
| `make_species_map.py` | 3 | $\geq 2$ species |
| `SOP-data2pair_list.py` | 4 | 1 species |
| `SOP-data2pair_list_multi.py` | 4 | $\geq 2$ species |
| `add_folded_bonds.py` | 5 | 1 species |
| `add_folded_bonds_multi.py` | 5 | $\geq 2$ species |
| `build_system.sh` | 1–5 | all systems |
| `ff_sop_42.input` | — | force-field file |
| `in_NVT.langevin` | — | NVT equilibration |
| `in_NPT.langevin` | — | NVT followed by NPT compression |

